# Buffering effects of nonspecifically DNA-bound RNA polymerases in bacteria

**DOI:** 10.1101/2023.11.04.565427

**Authors:** Yichen Yan, Tairan Li, Jie Lin

## Abstract

RNA polymerase (RNAP) is the workhorse of bacterial gene expression, transcribing rRNA and mRNA. Experiments found that a significant fraction of RNAPs in bacteria are nonspecifically bound to DNA, which is puzzling as these idle RNAPs could have produced more RNAs. Whether nonspecifically DNA-bound RNAPs have any function or are merely a consequence of passive interaction between RNAP and DNA is unclear. In this work, we propose that nonspecifically DNA-bound RNAPs buffer the free RNAP concentration and mitigate the crosstalk between rRNA and mRNA transcription. We verify our theory using mean-field models and an agent-based model of transcription, showing that the buffering effects are robust against the interaction between RNAPs and sigma factors and the spatial fluctuation and temporal noise of RNAP concentration. We analyze the relevant parameters of *Escherichia coli* and find that the buffering effects are significant across different growth rates at a low cost, suggesting that nonspecifically DNA-bound RNAPs are evolutionarily advantageous.

## I. INTRODUCTION

RNA polymerases (RNAPs) and ribosomes are finite resources that limit the overall rate of gene expression [1–11]. As a result, genes compete for these resources, leading to inevitable crosstalk of gene expression [12– 14]. This competition is particularly significant in bacteria as all RNAs are transcribed using the same type of RNAP. Thus, increasing the transcription of rRNA will inevitably reduce the availability of RNAPs for transcribing mRNA. However, despite this competition, a significant proportion of RNAPs in bacteria (30% to 50%) are nonspecifically bound to DNA and do not participate in transcription [15–19]. This phenomenon is puzzling as these idle RNAPs could have contributed to transcription, producing more mRNAs. Furthermore, producing these non-transcribing RNAPs may be a significant investment, and bacterial cells could have avoided producing these idle RNAPs and allocated the finite resources to other essential proteins to enhance their fitness. Indeed, eukaryotes have been found to suppress the nonspecific binding of RNAPs to DNA actively [20]. Therefore, a puzzle emerges: why do bacteria keep so many RNAPs nonspecifically bound to DNA?

In this study, we propose that nonspecifically DNA-bound RNAPs (which we refer to as nonspecific RNAPs in the following) in bacteria are crucial in regulating gene expression. These RNAPs mitigate the unwanted crosstalk between the expressions of different genes due to the limited availability of RNAPs. For example, when a subset of genes, such as the genes of rRNAs, are up-regulated, more RNAPs participate in their transcription, reducing the concentration of free RNAPs. This reduction in free RNAPs decreases the initiation rates of other genes since the probability of an RNAP binding to a promoter depends on the free RNAP concentration [15, 16, 18]. Nevertheless, nonspecific RNAPs help buffer the free RNAP concentration reduction and minimize the unwanted impact of resource competition (see the schematic in Fig. 1(a)). Similarly, when the expression of a group of genes is suppressed, nonspecific RNAPs buffer the increase in the free RNAP concentration. In this way, nonspecific RNAPs act like a pH buffer made of weak acid-base pairs, helping maintain the pH of an aqueous solution after adding an acid or base [21].

**FIG. 1.**
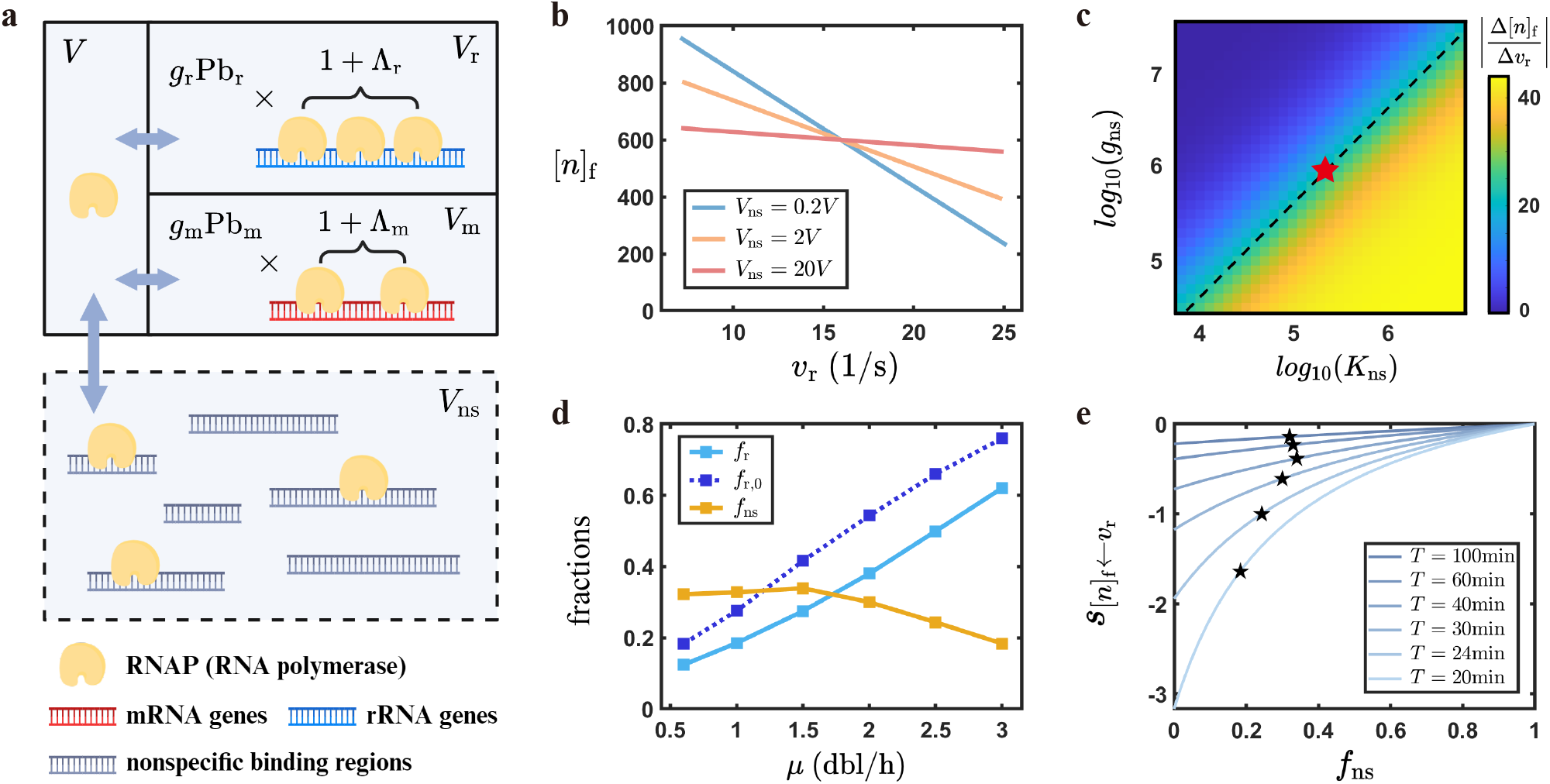
Nonspecific RNAPs buffer the free RNAP concentration upon gene expression changes. (a) The schematic diagram of the mean-field model. The nonspecific binding sites effectively enlarge the nucleoid volume, building a reservoir that exchanges RNAPs with the free RNAP pool. (b) The free RNAP concentration (unit: *µ*m^*−*3^) vs. the rRNA production rates under different effective volumes of nonspecific binding. The free RNAP concentration before perturbation under different *V*_ns_ is fixed at [*n*]_f_ = 600 *µ*m^*−*3^ (Table S1). The *V*_ns_ = 2*V* line corresponds to the parameters of *E. coli*. (c) The absolute slopes of [*n*]_f_ over *v*_r_ vs. *K*_ns_ (unit: *µ*m^*−*3^) and *g*_ns_. The slopes are equal as long as *g*_ns_*/K*_ns_ is the same (indicated by the dashed line with slope 1 in the logarithmic coordinates). The red star marks the corresponding values of *E. coli* at the 30-min doubling time (Table S1). (d) The estimated fractions of RNAPs on rRNA genes (*f*_r_), RNAPs on rRNA genes if the nonspecific binding is absent (*f*_r,0_), and nonspecific RNAPs (*f*_ns_) vs. growth rate for *E. coli*. (e) The sensitivity of free RNAP concentration to the changes in rRNA production rates as a function of the nonspecific RNAP fraction under different growth rates. Here, *T* is the doubling time, the inverse of the growth rate. The stars mark the *f*_ns_ values of *E. coli* and the corresponding y-values.

To assess the impact of nonspecific RNAPs on gene expression, we use a mean-field model of transcription neglecting spatial information [15, 16]. RNAPs can bind to promoters and nonspecific binding sites on DNA, and the probabilities of RNAP binding to these sites depend on the concentration of free RNAPs. Using analytical and numerical calculations, we demonstrate that non-specific RNAPs act as buffers for changes in free RNAP concentration caused by the regulation of specific genes, such as genes of rRNAs. The buffering effects help suppress crosstalk between different genes, promoting robust and stable gene expression. In particular, the physiological parameters of *Escherichia coli* across different growth rates are in the parameter space where the buffering benefit of nonspecific RNAPs is significant. In contrast, the cost of producing extra RNAPs is relatively low, suggesting that an appropriate amount of nonspecific RNAPs may be under evolution selection. We further extend our mean-field model by including sigma factors such that RNAP can bind to promoters and initiate transcription only if bound by a sigma factor. Our conclusions on the buffering effects of nonspecific RNAPs remain robust against the interaction between RNAPs and sigma factors. In particular, we find that if a cell aims to up-regulate a gene without affecting other genes, a good strategy is to change its promoter-RNAP dissociation constant; if a cell aims to up-regulate a group of genes and automatically suppress the others, a good strategy is to use sigma factor competition.

To strengthen our hypothesis, we also develop an agent-based model that explicitly incorporates spatial information, including the diffusion of RNA polymerases and their interactions with promoters and nonspecific binding sites. Importantly, the agent-based model naturally generates both spatial and temporal noise in the free RNAP concentration, from which we find that non-specific RNAPs indeed suppress the correlation between the production rates of mRNA genes and rRNA genes, thereby providing further evidence supporting our predictions.

On the application side, our findings suggest a strategy for reducing the interference between the expressions of exogenous and endogenous genes in synthetic biology [22, 23] by adding non-coding DNA sequences to the bacterial chromosome to increase the number of nonspecific binding sites.

## II. RESULTS

### A. Mean-field model

In the mean-field model, the probability of a promoter bound by an RNAP is a Hill function of the free RNAP concentration [15, 16, 18],

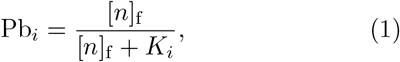

where [*n*]_f_ is the concentration of free RNAPs, and *K*_*i*_ is the dissociation constant of the promoter *i* (Methods A). In the mean-field model, we model gene expression regulation by changing the dissociation constant *K*_*i*_. The smaller *K*_*i*_ is, the stronger the promoter’s binding affinity to RNAP. Once an RNAP binds to a promoter, it transitions to the elongating state with an initiation rate 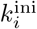, moves along the operon, and eventually detaches from DNA after it finishes transcribing. The number of elongating RNAPs (*n*_el,i_) on the operon following a promoter is proportional to the probability of the promoter bound by an RNAP:

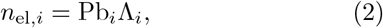

where 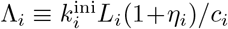, the maximum possible number of elongating RNAPs on the operon, which we name as capacity in the following. Here, *L*_*i*_ is the length of the operon following the promoter *i*, and *c*_*i*_ is the elongation speed of RNAP. When the number of RNAPs on the operon reaches the steady state, the number of initiations per unit time must be equal to the number of RNAPs that detach from the end of the operon, that is, 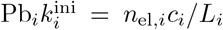, leading to Eq. (2) with *η*_*i*_ = 0. Here, we also include the pausing effect, that is, elongating RNAPs may pause for a finite duration, effectively increasing the capacity [18, 24]. *η*_*i*_ is the ratio between the number of pausing RNAPs and actively elongating RNAPs (Table S1).

We categorize RNAPs into three types: (1) promoter-bound RNAPs and elongating RNAPs (including pausing RNAPs), (2) free RNAPs diffusing within the nucleoid, (3) and nonspecifically DNA-bound RNAPs (i.e., nonspecific RNAPs) [15–18]. We consider a bacterial cell with a fixed nucleoid volume *V* and a fixed total number of RNAPs *n*. This simplification is reasonable as the time required for RNAPs to transcribe an operon (around one minute) is much shorter than the typical doubling time of bacteria (around 30 minutes or longer) [25]. Therefore, RNAP allocation to the different categories can be considered instantaneous. We use the nucleoid volume to compute the concentration because RNAPs are almost entirely within the nucleoid [26–28].

In this work, we aim to explore the buffering effects of nonspecific RNAPs such that specific genetic details are not critical to our conclusions. Therefore, we employ a coarse-grained approach by dividing the genome into two parts: promoters of rRNA and mRNA genes, each with their respective numbers (*g*_r_ and *g*_m_), dissociation constants (*K*_r_ and *K*_m_), and capacities (Λ_r_ and Λ_m_). The conservation of the total RNAP copy number *n*_t_ leads to the following equation:

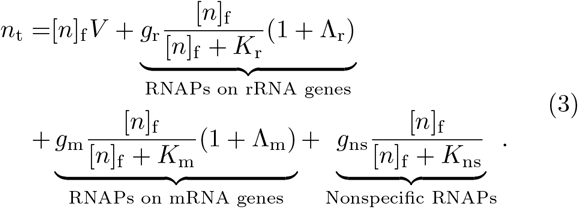

On the right side of the above equation, the last three terms represent the number of RNAPs binding to rRNA genes, mRNA genes, and nonspecific binding sites, respectively. The 1 and Λ_r,m_ in the 1 + Λ_r,m_ term correspond to promoter-bound RNAPs and elongating RNAPs. *g*_ns_ refers to the number of nonspecific binding sites on DNA, and *K*_ns_ represents the dissociation constant of nonspecific binding. RNAPs nonspecifically bound to DNA do not elongate along DNA; therefore, there is no extra capacity in the nonspecific RNAPs term of Eq. (3). Eq. (3) allows us to determine the free RNAPs concentration self-consistently.

To simplify the model, we introduce the idea of effective volumes and rewrite Eq. (3) as

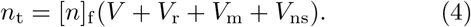

Here, 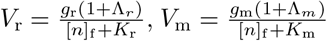, and 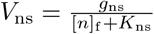.One can think of them as effective volumes that store DNA-bound RNAPs. Nonspecific binding sites equivalently enlarge the nucleoid volume, building a reservoir of nonspecific RNAPs (Fig. 1(a)). Finally, we define the RNA production rate as the number of RNAs produced per unit time,

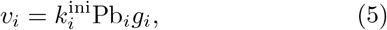

where *i* represents mRNA or rRNA genes.

### B. The buffering effects of nonspecific RNAPs on free RNAP concentration

#### 1. The buffering effects are significant in E. coli

We estimate the relevant parameters of the model organism, *E. coli* with a doubling time of 30 minutes (see Table S1 with the detailed calculations and references in-cluded in the caption). As expected, the promoter of the mRNA genes is much weaker than that of rRNA genes, and the binding affinity of nonspecific binding sites is even weaker. Meanwhile, the free RNAP concentration in this condition is about [*n*]_f_ = 600 *µ*m^*−*3^, suggesting that the rRNA promoter has a 1/3 probability of being occupied while the binding probability of mRNA promoter on average is much less than 1 (Eq. (1)).

We next study the impact of a sudden rRNA regulation on the free RNAP concentration by changing the dissociation constant of the rRNA promoter *K*_r_ from its unregulated value. Interestingly, using the parameters of *E. coli* (Table S1), the free RNAP concentration is weakly dependent on the rRNA production rate (the *V*_ns_ = 2*V* line in Fig. 1(b)). If we take a smaller effective volume of nonspecific RNAPs, *V*_ns_, with the same [*n*]_f_ before regulation, [*n*]_f_ becomes much more sensitive to the rRNA production rate. In contrast, if we make the *V*_ns_ larger, [*n*]_f_ can be almost independent of the rRNA production rate. Notably, one can change *V*_ns_ either by changing *K*_ns_ or changing *g*_ns_, and the results are the same as long as they lead to the same *V*_ns_ (see detailed proof in Methods B). Moreover, because [*n*]_f_ ≪ *K*_ns_ (Table S1) in typical biological scenarios, *V*_ns_ can be well approximated by *g*_ns_*/K*_ns_; therefore, the sensitivity is virtually the same as long as *g*_ns_*/K*_ns_ is the same (Fig. 1(c)). The buffering effects of nonspecific RNAPs are equivalently applicable to a sudden mRNA regulation (Fig. S1(a-b)).

#### 2. The buffering effects are parameter-insensitive and valid across growth rates

We define the sensitivity of [*n*]_f_ against the change in the rRNA production rate due to a small change in *K*_r_ as 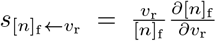. Surprisingly, we find that 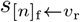 only depends on the fractions of different types of RNAPs,

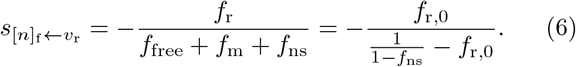

Here, *f*_free_, *f*_r_, *f*_m_, and *f*_ns_ represent the fractions of free RNAPs, RNAPs bound to rRNA genes, mRNA genes, and nonspecific binding sites (see their values for *E. coli* at the doubling time 30 min in Table S1). Here, *f*_r,0_ represents the fraction of RNAPs bound to rRNA genes if there is no nonspecific binding, which satisfies *f*_r,0_*/f*_r_ = 1*/*(1 −*f*_ns_). We note that *f*_ns_ and *f*_r,0_ are growth-rate dependent (Fig. 1(d), see Methods C for the details of calculations.)

Given a doubling time with its estimated *f*_r,0_, we tune the fraction of nonspecific RNAPs. Interestingly, for *E. coli*, the sensitivity of free RNAP concentration to the change in rRNA gene expression is about half compared to the case where nonspecific RNAPs are absent (Fig 1(e)). This result suggests that the endogenous numbers of nonspecific binding sites and their dissociation constant significantly buffer the change of the free RNAP concentration across different growth rates. Similarly, nonspecific RNAPs significantly reduce the sensitivity of free RNAP concentration to the changes in mRNA gene expression (Fig. S1(c)) with an equation similar to Eq. (6) holds as well (Eq. (15) in Methods B).

### C. The cost and benefit of nonspecific RNAPs

Although nonspecific RNAPs can attenuate unwanted crosstalk between genes, they also require the cell to spend additional resources to establish the reservoir of nonspecific RNAPs. To demonstrate whether nonspecific RNAPs are evolutionarily advantageous, we compare the cost and benefit of nonspecific RNAPs. To quantify the cost, we use the fraction of nonspecific RNAPs (which are considered as excess resources) in the entire proteome, *Φ*_ex_ = *Φ*_n_*f*_ns_ where *Φ*_n_ is the fraction of total RNAPs in the proteome. We compute its value for *E. coli* for different growth rates and find that they are typically small (Fig. 2(a), and see the details of calculation in Methods C).

**FIG. 2.**
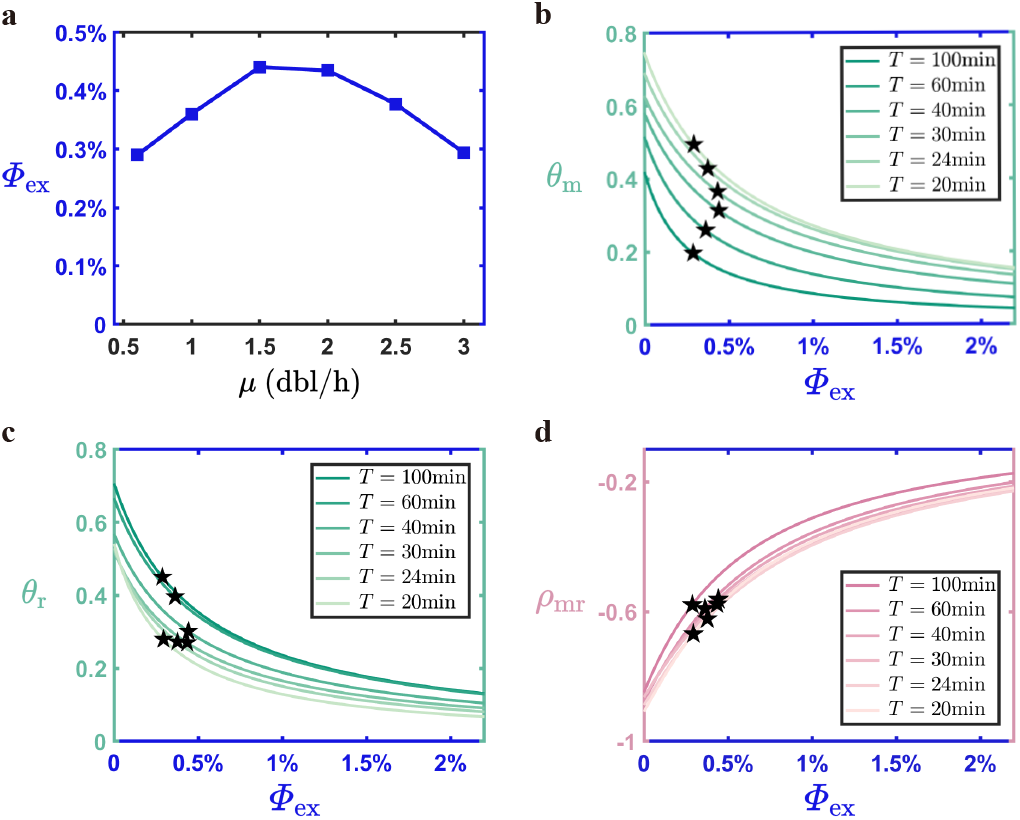
The cost and benefit of nonspecific RNAPs. (a) The cost of nonspecific RNAPs under different growth rates, quantified by the proteome fraction of nonspecific RNAPs *Φ*_ex_. (b-c) The crosstalk factors for mRNA genes (b) and rRNA genes (c) as a function of *Φ*_ex_ under six different growth rates. Here, *T* is the doubling time, the inverse of the growth rate. The stars mark the *Φ*_ex_ values of *E. coli* and the corresponding y-values. (d) The correlation coefficient of mRNA and rRNA production rates as a function of the proteome fraction of nonspecific RNAPs. The stars mark the values of *E. coli*. The detailed calculations of *Φ*_ex_, *θ*_m_, *θ*_r_ and *ρ*_mr_ are shown in Methods C-E.

To quantify the benefit, we introduce perturbations to the dissociation constants *K*_m_ and *K*_r_ and calculate the relative changes of mRNA and rRNA production rate through the following sensitivity matrix (Methods D):

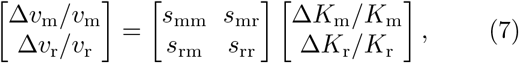

where *s*_mm_ and *s*_rr_ are the self-sensitivity factors, and *s*_mr_ and *s*_rm_ are the mutual-sensitivity factors. We quantify the benefit as the crosstalk factors: *θ*_m_ = |*s*_mr_*/s*_mm_| and *θ*_r_ = |*s*_rm_*/s*_rr_|, which quantify the degree of gene expression crosstalk (Methods D).

Interestingly, we find that for *E. coli*, the crosstalk factors of mRNA and rRNA are approximately reduced by half with the cost of a small nonspecific RNAP fraction in the proteome across different growth rates (Fig. 2(b-c)). This result demonstrates that the buffering effects of nonspecific RNAPs do not incur a significant cost, suggesting that they are likely the consequence of evolution selection. We also introduce another quantitative cost trait: the excess factor *ε*, defined as the number of non-specific RNAPs divided by the number of other RNAPs: 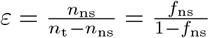, and our conclusions remain the same (Fig. S3).

We also study the buffering effects against random gene production rates to mimic a bacterial cell exposed to a fluctuating environment. We assume that the dissociation constants fluctuate with the following noise levels: 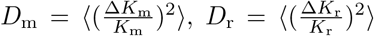. Using Eq. (7), the Pearson correlation coefficient of the mRNA production rate and rRNA production rate is analytically calculated [Eq. (21)]. Notably, the values of *Φ*_ex_ of *E. coli* across different growth rates are again within the range where the buffering against temporal noise is significant with a small cost, showing the robustness of our results (Fig. 2(f)). Here, we use *D*_m_ = *D*_r_ = 0.2, and our conclusions remain the same under different values of *D*_m_ and *D*_r_ (Fig. S4).

### D. Effects of sigma factors

#### 1. The RNAP partitioning rules considering sigma factors

In bacteria, sigma factors bind RNAPs and are required for RNAPs to recognize promoters. RNAPs without sigma factor binding are called core RNAPs, and core RNAPs become holoenzymes only if they are bound by a sigma factor [29–31]. While both core RNAPs and holoenzymes nonspecifically bind to DNA, only holoenzymes specifically bind to promoters and initiate transcription. Meanwhile, anti-sigma factors bind to sigma factors and prevent their binding with RNAPs, inhibiting transcription [31–34]. It is unknown whether the buffering effects of nonspecific RNAPs are still valid in the presence of multiple types of sigma factors, and the existence of anti-sigma factors makes the conclusions even more unclear. To test the robustness of our results, we extend our model to include sigma and anti-sigma factors (Methods F). We use *σ*^*j*^ to represent the type *j* sigma factor with its corresponding anti-sigma factors *Anti*^*j*^, *E* to represent the core RNAP, and *Eσ*^*j*^ to represent the corresponding holoenzyme. The total RNAP number as the sum of core RNAPs and all types of holoenzymes is still represented by *n*_t_. The total free RNAP concentration as the sum of free core RNAP concentration ([*E*]_f_) and all types of free holoenzyme concentration ([*Eσ*^*j*^]_f_) is still represented by [*n*]_f_ (Eq. (29)).

In this extended model with sigma factors, we assume that the transcription of each gene is initiated by a particular type of holoenzyme, which becomes a core RNAP during elongation because the sigma factor quickly dissociates from the holoenzyme early after transcription initiation [35–38]. Therefore, the RNA production rate of gene *i* is determined by its corresponding free holoenzyme concentration [*Eσ*^*j*^]_f_ :

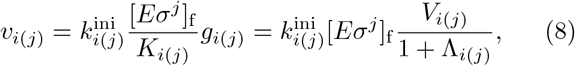

where *i*(*j*) denotes gene *i* recognized by *σ*^*j*^. 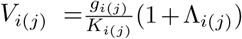 is the effective volume of gene *i* that store holoenzymes on promoter *i* and the subsequent elongating core RNAPs, that is, [*Eσ*^*j*^]_f_ *V*_*i*(*j*)_ is the total number of RNAPs on gene *i*. We have used the approximation 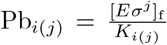 (Eq. (1)), which is biologically reasonable since the probability of a promoter bound by a holoenzyme is typically low (Fig. S2(b)). Moreover, this approximation significantly simplifies the analytical derivations and numerical simulations (Methods F). Similarly, [*Eσ*^*j*^]_f_ *V*_ns_ and [*E*]_f_ *V*_ns_ are the numbers of holoenzymes and core RNAPs nonspecifically bound to DNA, respectively. Here, we approximate 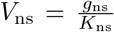 because *K*_ns_ ≫ [*Eσ*^*j*^]_f_, and *K*_ns_ ≫ [*E*]_f_ in typical biological scenarios (Table S1), where we assume core RNAPs and holoenzymes have the same *K*_ns_ because their nonspecific binding affinities with DNA are comparable [39].

Intriguingly, based on several equilibrium equations of chemical reactions and conservation equations (Fig. 3(a) and Methods F) we derive two partition equations. The first one tells us the fractions of different types of RNAPs in the total pool of RNAPs (Fig. 3(b), left):

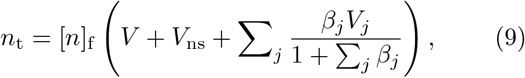

where [*n*]_f_ *V*, [*n*]_f_ *V*_ns_ and 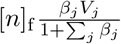 are the numbers of free RNAPs, nonspecific RNAPs, and RNAPs bound to genes recognized by *σ*^*j*^, respectively. Here, *V*_*j*_ = Σ_*i*(*j*)_ *V*_*i*(*j*)_ is sum of effective volumes over all genes recognized by *σ*^*j*^; *β*_*j*_ is the partition factor (see its detailed expression in Methods F), which is positively related to the total sigma factor number and negatively related to the total anti-sigma factors and holoenzyme dissociation constants. The second one tells us how the free RNAPs are partitioned into different types of free holoenzymes and free core RNAPs (Fig. 3(b), right):

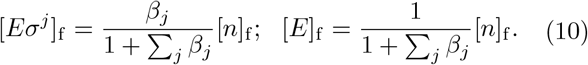

Eq. (9) and (10) together determine the concentration of a specific type of free holoenzymes and subsequently determine the RNA production rate of a particular gene.

**FIG. 3.**
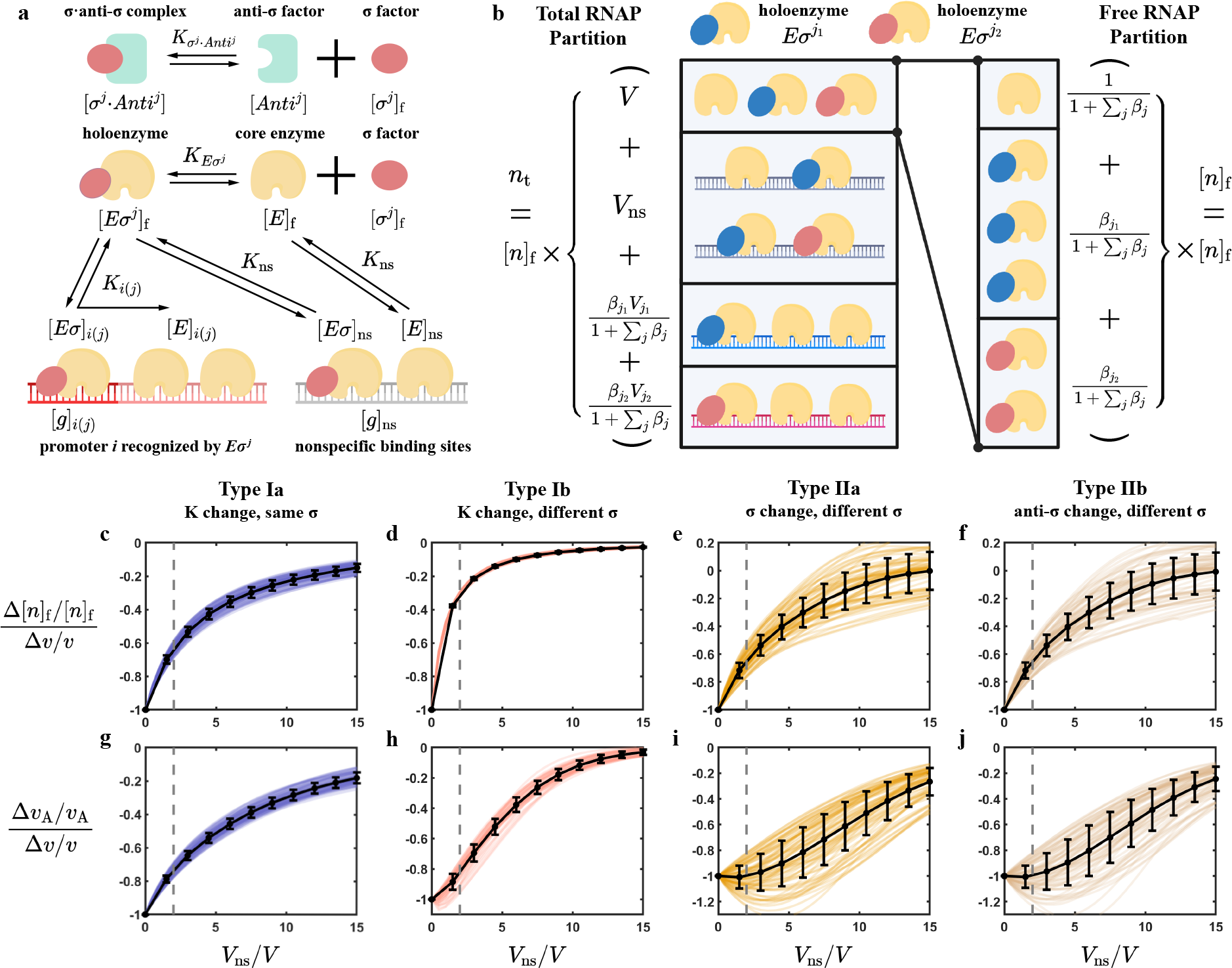
The buffering effects of nonspecific RNAPs in the model with sigma factors. (a) The schematic of the mean-field model with sigma and anti-sigma factors included. Both core RNAPs and holoenzymes nonspecifically bind to DNA with the same dissociation constant *K*_ns_. (b) The schematic of the partitioning of total RNAPs and free RNAPs based on Eq. (9) and Eq. (10). (c-f) Nonspecific RNAPs can buffer the free RNAP concentration regardless of the types of regulation. The y-axis shows the sensitivity of free RNAP concentration to the changes in the RNA production rate of the regulated gene, which can be gene B or C, depending on the regulation type. The values under different sets of parameters are scaled to *−*1 at *V*_ns_ = 0. The curves with random parameters are shown in the background, and the black lines are their means. The error bars are the standard deviation of the background curves. The dashed line marks the *V*_ns_*/V* value of *E. coli*, which is approximately constant across growth rates (Methods C). (g-j) The same as (c-f), but the y-axis shows the sensitivity of the RNA production rate of genes A to the changes in the RNA production rate of genes B or C, depending on the regulation type. Nonspecific RNAPs attenuate gene expression crosstalk significantly in type I (dissociation-constant regulation), but non-significantly in type II (sigma competition).

#### 2. The buffering effects of nonspecific RNAPs are mostly valid considering sigma factors

One should note that Eq. (9) is essentially identical to Eq. (4) where sigma factors are neglected, except that the effective volume of genes is multiplied by the factor 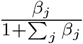. Therefore, our previous conclusion that the free RNAP concentration [*n*]_f_ (including both core RNAPs and holoenzymes) can be buffered by nonspecific RNAPs is still valid. However, a stable free RNAP concentration does not guarantee stable gene expression, as it is the free holoenzyme concentration that determines the RNA production rate. Therefore, we numerically solve Eq. (9) and (10) to illustrate the roles of nonspecific RNAPs.

We remark that the exact values of parameters, e.g., the copy numbers of sigma factors, are uncertain. For example, some experiments showed that the number of total sigma factors is more than the total RNAPs, while others showed the opposite [40–44]. The parameters under different conditions and growth rates are also distinct. Therefore, we randomly sample the parameters from an extensive range to ensure that our conclusions are independent of specific parameters’ values (Table S2).

Like the previous sections, we coarse-grain the genome into three representative “genes.” Genes A and B are under the control of sigma factor 1, and gene C is under the control of sigma factor 2. We study gene expression crosstalk by focusing on the expression level of gene A with four scenarios. (Ia) We tune the dissociation con-stant of gene B and see how it affects the expression of gene A. (Ib) We tune the dissociation constant of gene C and see how it affects the expression of gene A. (IIa) We tune the number of sigma factor 2 and see how it affects the expression of gene A. (IIb) We tune the number of anti-sigma factor 2 and see how it affects the expression of gene A. One should note that Case Ia is essentially what we study in the previous sections.

Interestingly, we find that the free RNAP concentration [*n*]_f_ is buffered by the nonspecific RNAPs in all four cases (Fig. 3(c-f)). When it comes to gene expression crosstalk, nonspecific RNAPs are still effective in reduc-ing crosstalk due to changes in the promoter-RNAP dissociation constant no matter whether the regulated gene is under the control of the same sigma factor as gene A or not (Fig. 3(g-h)). Nevertheless, nonspecific RNAPs are relatively weaker in buffering sigma competition; that is, when the expression level of gene C is tuned by the changes in the number of its sigma factor or anti-sigma factor, the expression level of gene A can be significantly affected even with a large reservoir of nonspecific RNAPs (Fig. 3(i-j)). This indicates that if the cell aims to up-regulate a gene without affecting other genes, a good choice is to change its promoter-RNAP dissociation constant; if the cell aims to up-regulate a group of genes and automatically suppress the others, a good option is by sigma factor competition. We also include a more mathematical explanation for this difference in the attenuation effects of nonspecific RNAPs on gene expression crosstalk in Methods F.

In the following section, we study an agent-based model with spatial information included. For simplicity, we focus on the scenario in which gene expression is regulated by the RNAP-promoter dissociation constant. We also neglect sigma factors since the model without sigma factors is enough to demonstrate the buffering effects of nonspecific RNAPs.

### E. Agent-based simulations

The mean-field model assumes that once an RNAP leaves a promoter or nonspecific binding site, it enters the well-mixed pool of free RNAPs. However, this model does not capture that the free RNAP concentration is spatially heterogeneous, and RNAPs that have just left a binding site are more likely to rebind to the same site. It also neglect the noise in the random processes during transcription. To test whether the buffering effects of nonspecific RNAP are still valid in the presence of the spatial effect and temporal noise, we develop an agent-based model which explicitly incorporates spatial information, including the diffusion of RNA polymerases and their interactions with promoters and nonspecific binding sites. Furthermore, the agent-based model naturally generates fluctuating mRNA and rRNA production rates, from which we can test whether nonspecific RNAPs suppress the correlation between the mRNA and rRNA gene expression robustly.

This model represents the nucleoid as a three-dimensional space with multiple binding sites corresponding to promoters and nonspecific binding sites. We model RNAPs as point particles, diffusing in the nucleoid with a diffusion constant *D* until they hit any free binding site (Fig. 4(a)). If RNAPs enter a free nonspecific binding site, they will leave with an off-rate of 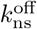. If RNAPs enter a promoter, they will either leave with an off-rate of 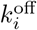 or transition to the elongation state with a rate of 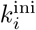 (Fig. 4(b)), where *i* is either “r” or “m”, depending on whether the promoter is followed by rRNA or mRNA genes [45]. Once a binding site is occupied, it cannot bind another RNAP until the RNAP hops off or transitions to the elongation state. An elongating RNAP becomes free after a fixed duration, which is the time for an RNAP to finish transcribing an operon, i.e.,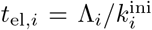 (Table S1). The rRNA (mRNA) production rate is computed as the number of elongating RNAPs on all rRNA (mRNA) genes divided by *t*_el,*i*_, which is the number of RNA produced per unit time because each elongating RNAP transcribes one RNA.

**FIG. 4.**
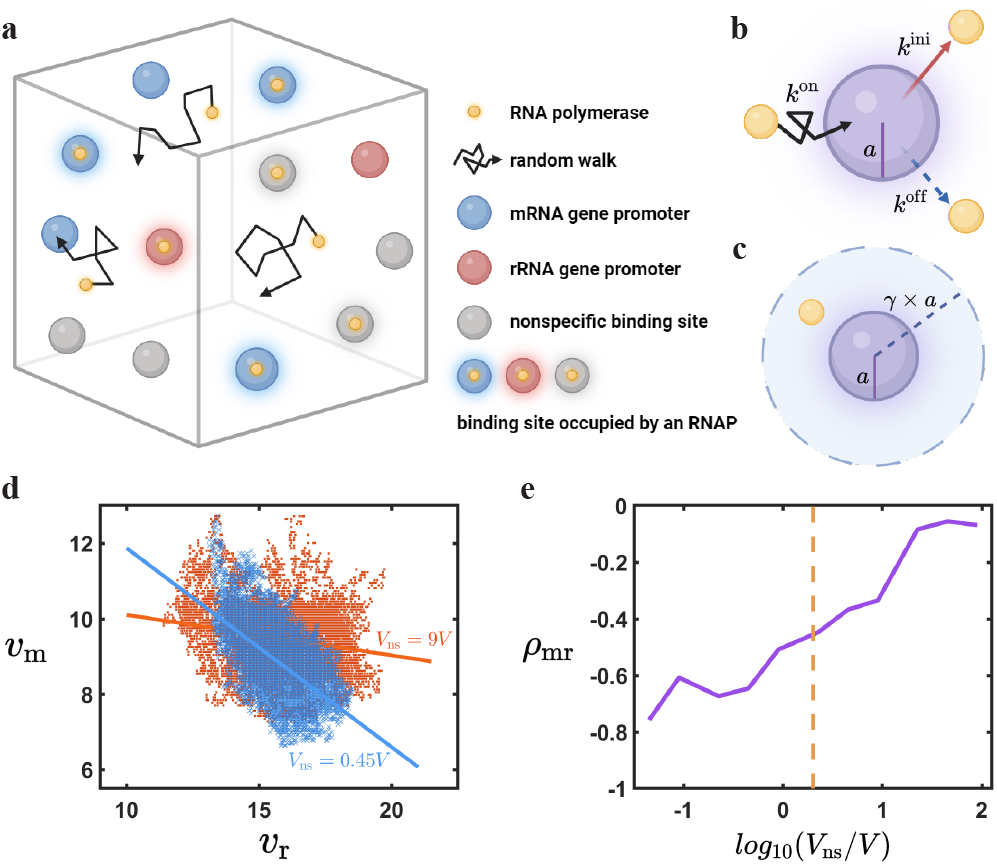
Simulations of the agent-based model. (a) A schematic of the agent-based model of bacterial transcription. We model the three-dimensional nucleoid with multiple binding sites corresponding to promoters and nonspecific binding sites. RNAPs diffuse until they hit any free binding site. (b) If an RNAP hits a promoter, it will start transcription with a rate 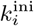 or hop off the promoter with a rate 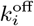. Here, *i* can be “r” or “m”, depending on whether rRNA or mRNA genes follow the promoter. If an RNAP hits a nonspecific binding site, it will only hop off with a rate 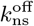. (c) We relocate an RNAP that just left a binding site inside a shell with inner diameter *a* and outer diameter *R* = *γ × a*. (d) The scatter plot of mRNA and rRNA production rates (unit: 1/s) under different *V*_ns_. The negative correlation between *v*_m_ and *v*_r_ across time is strong under weak nonspecific binding (small *V*_ns_) while weak under strong nonspecific binding (large *V*_ns_) (Methods H). Here, each point represents one time point in each repeat, and the straight line is the linear fit of all points of the corresponding *V*_ns_. (e) The absolute value of the negative correlation coefficient of mRNA and rRNA production rates decreases with the effective volume of nonspecific binding. The dashed line marks the *V*_ns_*/V* value of *E. coli*, which is approximately constant across growth rates (Methods C).

We represent the binding sites as spheres with radii *a*. Once an RNAP leaves a binding site, we relocate it randomly inside a shell with inner diameter *a* and outer diameter *R* = *γ* × *a* (Fig. 4(c)). Here, *γ >* 1 is the relocation parameter. For simplicity, we assume random and fixed positions of binding sites, and the binding of RNAPs to promoters and nonspecific binding sites are diffusion-limited. More details of the agent-based model are included in Methods G-H.

Intriguingly, the simulated free RNAP concentration (averaged over binding sites) vs. the distance to the binding site center can be well fitted by our analytical predictions (Fig. S5(a-b)), and the agent-based simulations agree well with the mean-field predictions regarding the sensitivity of average free RNAP concentration to the changes in gene expression (Fig. S5(c)) despite the spatial heterogeneity in the free RNAP concentration. We find a strong negative correlation between the mRNA and rRNA production rates given weak nonspecific binding (small *V*_ns_) and an almost zero correlation between them given strong nonspecific binding (large *V*_ns_) (Fig. 4(e-f)). Therefore, we conclude that the buffering effects of nonspecific RNAPs on gene expression crosstalk are robust against the spatial heterogeneity of free RNAP concentration and noises in random processes.

## III. DISCUSSION

Previous studies have shown that nonspecific transcription factor-DNA binding can buffer against gene expression noise in eukaryotes [46–48]. Similarly, it has been suggested that bacteria produce extra DnaA (master regulator of replication initiation) to suppress noise in initiation [49]. Recent studies highlight the importance of a trade-off between protein overabundance and cell fitness, especially for transcription machinery [50]. Nevertheless, the benefit compared to the cost of nonspecific binding has yet to be estimated as far as we realize, neither for transcription factors in eukaryotes nor RNAPs in bacteria. One should note that nonspecific binding of RNAPs appears absent in eukaryotes [20], which means that our conclusions mostly apply to bacteria.

In this work, we demonstrate that the nonspecifically DNA-bound RNAPs are natural buffers to mitigate the unwanted crosstalk between the transcription of rRNA and mRNA, and the buffering effects are significant in *E. coli*. Importantly, we mathematically prove that our conclusions are parameter-insensitive across different growth rates. We find that the buffering effects of nonspecific RNAPs are more significant against rRNA regulation in rapid-growth conditions (Fig. 1(e)). In contrast, the buffering effects are more significant against mRNA regulation in slow-growth conditions (Fig. S1(c)). We argue that this growth rate dependence is due to the growth rate dependence of the fractions of different types of RNAPs (Eqs. (14, 15), Fig. 1(d) and Fig. S2). We note that an additional benefit of nonspecific binding at slow growth can be providing a reservoir for transitions to fast growth with high transcription rates of rRNA.

Interestingly, the number of nonspecific binding sites is much more than that of promoters, and the nonspecific binding affinity is much weaker than those of promoters, similar to pH buffers made of weak acid-base pairs (Table S1). The buffering effects of nonspecific RNAPs to maintain a stable free RNAP concentration may help bacteria to maintain robust regulation of transcription initiation rates across the genome [11]. A stable mRNA level can be important for cell physiologies as it may influence protein production and the growth rate since ribosomes compete for transcripts [51].

Meanwhile, producing nonspecific RNAPs also has a cost because cells have to make more RNAPs to keep the same free RNAP concentration. We propose that the benefit of weaker interference between genes and the cost of producing extra RNAPs together set the number of nonspecific binding sites and their binding affinity through evolution. We cannot exclude the possibility that the origin of nonspecific binding may result from passive interaction between RNAP and DNA. But still, whether it is passive or selected actively, one cannot ignore the function of nonspecifically DNA-bound RNA polymerases as a buffer of resource competition.

In growing cells, the number of *σ*^70^ is significantly larger than other alternative types of sigma factors in *E. coli* [40, 41, 52], and most genes are controlled by *σ*^70^. However, in the presence of multiple types of sigma factors [41, 53–55], different types of sigma factors can compete for the pool of core RNAP such that the upregulation of one group of genes can result in reduced expressions of genes controlled by other types of sigma factors, which is called sigma competition. It has been suggested that nonspecific RNAPs have mild effects on mitigating the crosstalk between different groups of genes due to sigma competition [52]. Here, we show that this result does not contradict the buffering effects between genes controlled by the same type of sigma factor. Unexpectedly, we find that nonspecific RNAPs help reduce the crosstalk between genes regardless of whether they are regulated by the same type of sigma factors, as long as gene regulation is implemented through the RNAP-promoter dissociation constants. Meanwhile, cells can use alternative sigma factors to up-regulate some genes while down-regulating others through the changes of sigma factor and anti-sigma factor copy numbers.

Our model neglects the spatial correlation of promoter locations while spatial and temporal organizations of the nucleoid have been observed [19, 56–58]. In the future, it will be interesting to study how the nucleoid structure influences our conclusions. Furthermore, it is also essential to verify whether the buffering effects due to nonspecifically DNA-bound RNAPs apply to other bacteria. On the application side, our work provides a new avenue to reduce the burden effects of synthetic circuits on bacteria by adding non-coding DNA to the chromosome to increase the number of nonspecific binding sites.

## VI. METHODS

### A. The probability of a promoter or a nonspecific binding site bound by an RNAP

We assume that the binding of RNAPs to promoters is diffusion-limited. Furthermore, once an RNAP binds a promoter, it will either leave with an off-rate 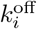 or transition to the elongation state with a rate 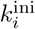. The probability of promoter *i* bound by an RNAP (Pb_*i*_) is determined by the equilibrium between the formation and dissociation of the transcription complex: 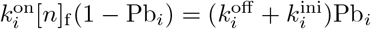, which gives

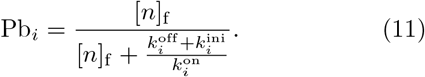

Comparing Eq. (11) to Eq. (1), we have 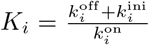. For nonspecific binding sites, there is no elongation; therefore, 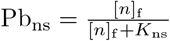 with 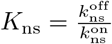.

### B. The sensitivity of [*n*]_f_ to the changes in rRNA (mRNA) production rates

We replace the number of RNAPs on rRNA genes term ([*n*]_f_ *V*_r_) in Eq. (4) by the rRNA production rate *v*_r_ such that

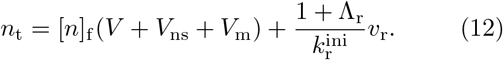

Considering a small change in *K*_r_ (notice that *V*_ns_ and *V*_m_ are functions of [*n*]_f_), we get 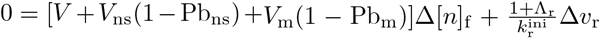. Rearranging the above equation and using the approximation Pb_ns_ ≪ 1, Pb_m_≪ 1, which is biologically reasonable (Eq. (1) and Table S1), we obtain

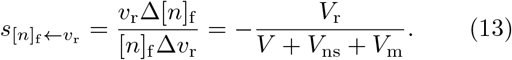

Eq. (13) expresses 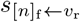 as a function of *V*_ns_. Because in typical biological scenarios, [*n*]_f_ ≪ *K*_ns_ (Table S1), we have *V*_ns_ = *g*_ns_*/K*_ns_, which means that changing *K*_ns_ or changing *g*_ns_ leads to the same buffering effects as long as they lead to the same *g*_ns_*/K*_ns_. We further rewrite Eq. (13) as

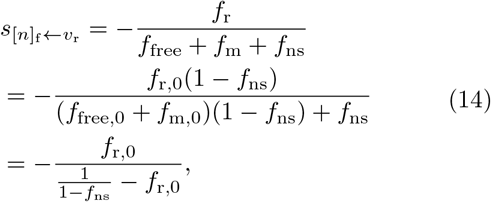

which is Eq. (6). Here, *f*_free,0_, *f*_m,0_ and *f*_r,0_ represent the fractions of free RNAPs and RNAPs bound to mRNA genes and rRNA genes if there is no nonspecific binding, which satisfy *f*_free_*/f*_free,0_ = *f*_m_*/f*_m,0_ = *f*_r_*/f*_r,0_ = 1 − *f*_ns_, respectively. The relationship *f*_free,0_ + *f*_m,0_ = 1 − *f*_r,0_ is used to get the last line. Similarly, we can get

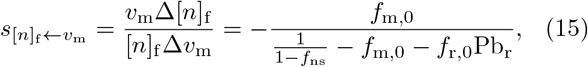

which is shown in Fig. S1(c). Here, Pb_r_ is not neglected as its value is comparable to other terms (Fig. S2).

### C. The growth-rate-dependent parameters for *E. coli*

We first determine the fractions of different types of RNAPs at the 30-min doubling time. *f*_free_, *f*_ns_, and (*f*_m_ + *f*_r_) at the 30-min doubling time are measured by single-molecule tracking due to the differences in the diffusion coefficients among free RNAPs, nonspecific RNAPs, and specifically-bound RNAPs [17]. For simplicity, we neglect a small fraction of RNAPs (about 0.06) unable to bind to DNA, which is likely immature RNAPs [16, 18]. To ensure that *f*_free_ + *f*_r_ + *f*_m_ + *f*_ns_ = 1, the fractions we estimate are slightly larger than the reported mean values but still within the ranges of experimental errors [17], which are *f*_free_ = 0.15, *f*_ns_ = 0.3, *f*_m_ + *f*_r_ = 0.55. The ratio of actively elongating RNAPs transcribing rRNA to those transcribing mRNA is about 2.2 [59]; therefore, we obtain *f*_r_ = 0.38 and *f*_m_ = 0.17. The ratio between pausing RNAPs and actively elongating RNAPs for rRNA genes and mRNA genes together is (0.54 − 0.22)*/*0.22 = 1.5 [17, 18, 59] (Details in Table S1(9)). For simplicity, we assume the ratios for rRNA and mRNA genes are the same and equal to 1.5.

Next, we determine the RNAP fractions across different growth rates. We calculate the fractions of actively elongating RNAPs on mRNA and rRNA genes across dif-ferent growth rates (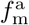 and 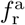) using the total RNAP number and the number of active RNAPs transcribing mRNA and rRNA at different growth rates in Ref. [59]. The ratio between pausing RNAPs and actively elongating RNAPs on mRNA and rRNA genes (*η*_m_ and *η*_r_) are negatively related to growth rates due to ppGpp regulation [24, 60]. For simplicity, we assume that *η*_m_ and *η*_r_ are the same, and both linearly decrease with growth rates and approach 0 when the doubling time *T* = 12 min. This leads to *η*_r_ = *η*_m_ = 1.5 − 0.5(*µ* − 2), and the detailed choices of *η*_r_ and *η*_m_ do not affect our main conclusions (Fig. S7). We then obtain 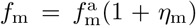 and 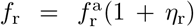. The ratio of *f*_ns_ to *f*_free_ is *V*_ns_*/V* = (*g*_ns_*/V*)*/K*_ns_ since *K*_ns_ ≫ [*n*]_f_, which is approximately constant across growth rates given a constant density of DNA and a constant *K*_ns_. Thus, we set this constant ratio as 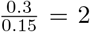 using the data of the 30-min doubling time and we obtain 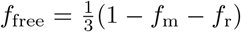 and 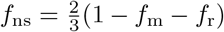.*f*_free,0,_*f*_m,0_ and *f*_r,0_ are computed from *f*_free_*/f*_free,0_ = *f*_m_*/f*_m,0_ = *f*_r_*/f*_r,0_ = 1 − *f*_ns_. *f*_r_, *f*_r,0_ and *f*_ns_ are shown in Fig. 1(d), and the other fractions are shown in Fig. S2(a).

We estimate the probabilities of the promoter bound by an RNAP for mRNA genes and rRNA genes from the fractions of actively elongating RNAPs: 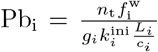 (Fig. S2(b)), where *i* represents mRNA or rRNA genes. The maximum transcription initiation rate 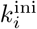 and gene length *L*_*i*_ are growth-rate independent [16, 18, 61] (Table S1). *n*_t_, 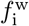, *g*_*i*_, and *c*_*i*_ are growth-rate dependent, and their values are from Ref. [59]. The two indexes of cost of nonspecific binding are calculated by: *Φ*_ex_ = *Φ*_n_*f*_ns_ (Fig. 2(a)) and 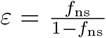 (Fig. S3(a)), where *Φ*_n_ is the growth-rate-dependent RNAP proteome fraction [59, 62].

### D. The sensitivity matrix and crosstalk factors

We consider a small change in both *K*_r_ and *K*_m_, and using Eq. (4), we obtain the change in the free RNAP concentration:

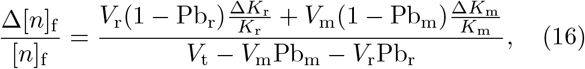

where *V*_t_ = *V* + *V*_ns_ + *V*_m_ + *V*_r_. Similarly, for the mRNA production rate 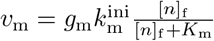, we have

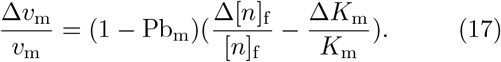

Combing Eq. (16) and Eq. (17) and using the approximation Pb_m_ ≪ 1 (Fig. S2(b)), we have

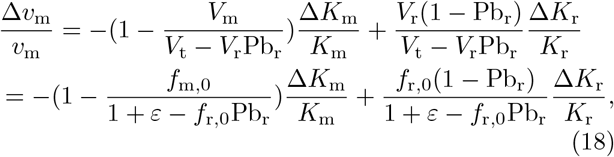

where we introduce the excess factor 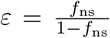. Simi-larly, for the rRNA production rate we have

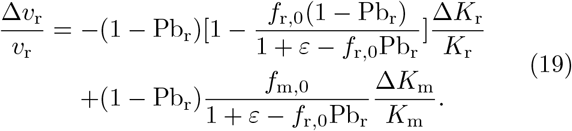

Writing the above two equations in matrix form, we finally get Eq. (7), and the sensitivity matrix is:

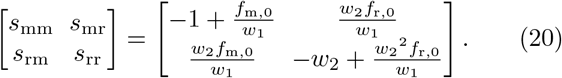

Here, *w*_1_ = 1 + *ε* − *f*_r,0_Pb_r_, and *w*_2_ = 1 − Pb_r_. Notice that *w*_1_ increases with *ε* linearly, so with the increase of nonspecific binding, the self-sensitivity factors (*s*_mm_ and *s*_rr_) approach their maximum absolute values, while the mutual sensitivity factors (*s*_mr_ and *s*_rm_) decrease to 0.

We define the crosstalk factors for mRNA and rRNA genes as *θ*_m_ = |*s*_mr_*/s*_mm_| and *θ*_r_ = |*s*_rm_*/s*_rr_|, and compute their values using the fractions of different types of RNAPs and probabilities of promoter binding estimated in section C of Methods.

### E. The correlation coefficient between the mRNA and rRNA production rates

We compute the covariance of *v*_m_ and *v*_r_ as Cov(*v*_m_, *v*_r_) = (*s*_mm_*s*_rm_*D*_m_ + *s*_rr_*s*_mr_*D*_r_)*v*_m_*v*_r_, and the product of the variances of *v*_m_ and *v*_r_ as 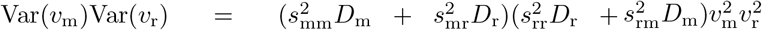. Here, Eq. (7) and ⟨Δ*K*_r_Δ*K*_m_⟩ = 0 are used. Finally, we get the correlation coefficient between mRNA and rRNA production as

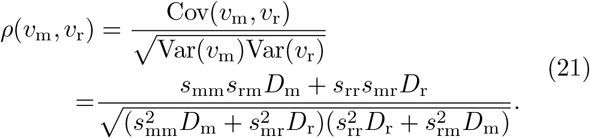

### F. Mean-field model including sigma factors

The concentration of holoenzymes on promoter *i* is related to the corresponding free holoenzyme concentration by:

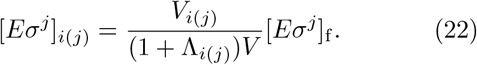

Similarly, the concentrations of core RNAPs and holoenzymes on nonspecific binding sites are:

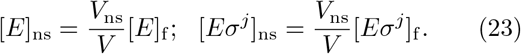

The following equation describes the equilibrium condition of sigma-core binding and holoenzyme dissociation:

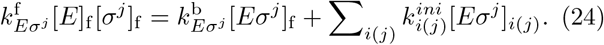

Here, [*σ*^*j*^]_f_ is the concentration of free sigma factors unbound by anti-sigma factors or core RNAPs. [*E*]_f_ is the concentration of free core RNAPs, and [*Eσ*^*j*^]_f_ is the concentration of free holoenzymes. 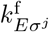 and 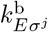 are the forward and backward rates of holoenzyme formation. The second term on the right side of the equation comes from the dissociation of holoenzymes into core RNAPs and sigma factors after transcription initiation. Therefore, the dissociation constant of core RNAP-sigma bind-ing becomes:

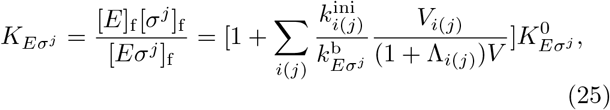

where 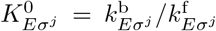 is the standard dissociation constant without transcription.

Next, we write the equilibrium equation for the sigmaanti-sigma complex and the conservation equation for anti-sigma factors:

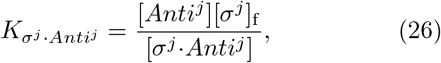

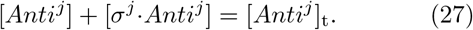

Here, [*Anti*^*j*^] is the concentration of anti-sigma factors without binding to sigma factors, [*σ*^*j*^·*Anti*^*j*^] is the concentration of sigma-anti-sigma complexes, and [*Anti*^*j*^]_t_ is the total anti-*σ*^*j*^ concentration.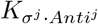 is the corresponding dissociation constant.

For each type of sigma factor, the concentrations of free sigma factors, sigma-anti-sigma complexes, free holoenzymes, and holoenzymes bound to DNA add up to the concentration of total sigma factors:

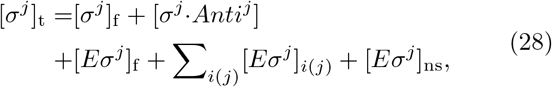

where [*σ*^*j*^]_t_ is the concentration of total sigma factors *j*. Here, we assume that the sigma factor quickly dissociates from the holoenzyme early after transcription initiation [35–38]. Therefore, the above sum does not include elongating RNAPs.

The concentrations of free core RNAPs and all free holoenzymes add up to the concentration of all free RNAPs, [*n*]_f_ :

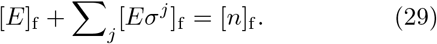

Combining Eqs. (22-23,25-29) leads to the following two free RNAP partition equations:

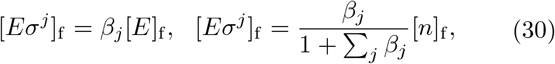

which are Eq. (10). Here, 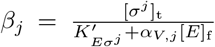 is the partition factor with 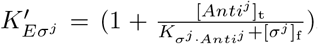 and 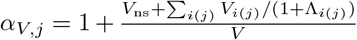. The expression of 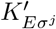 implies that anti-sigma factors make the dissocia-tion constant 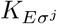 effectively larger. One should note that it is the linear combination of *V*_ns_ and *V*_*i*(*j*)_ that determines *α*_*V,j*_ and *β*_*j*_, which explains why nonspecific RNAPs are more effective in attenuating the gene ex-pression crosstalk when the regulation is implemented by changes in the dissociation constants. At last, the conservation equation for all RNAPs is

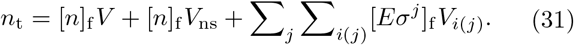

Replacing [*Eσ*^*j*^]_f_ by Eq. (30), we obtain Eq. (9).

### G. The free RNAP concentration distribution and *k*^on^ in the agent-based model

In the agent-based model, the free RNAP concentration [*n*]_f_ (*r, t*) becomes spatially dependent due to the absorption and release of RNAPs by the binding sites, which satisfies the following generalized diffusion equation,

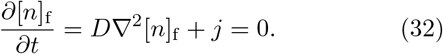

Here, we assume that once RNAPs leave binding sites, they are randomly relocated inside a shell around the center of the binding site with inner diameter *a* and outer diameter *R* = *γ* × *a* and *j* is the RNAP relocation flux per unit volume. For *r > R, j* = 0. The solution of Eq. (32) is

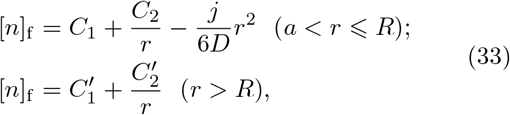

where *C*_1_, *C*_2_, 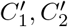 are constants defined by boundary conditions, which are

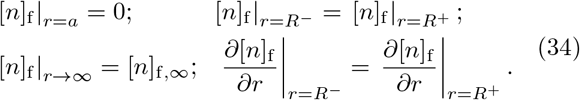

Here, [*n*]_f,∞_ is the asymptotic free RNAP concentration. These lead to the solutions (notice *γ* = *R/a*)

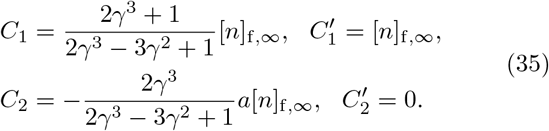

In the steady state, the relocation flux density *j* is related to the total flux entering a binding site *J* through

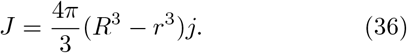

Meanwhile, *J* is also proportional to the gradient of the free RNAP concentration near the binding site surface, 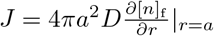. So from Eq. (33), we obtain

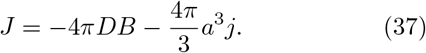

Combining Eq. (35-37), we get

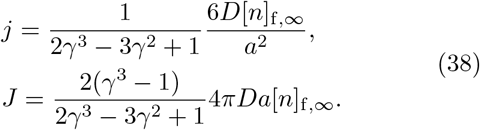

Finally, according to the definition *k*^on^ = *J/*[*n*]_f,∞_, we obtain:

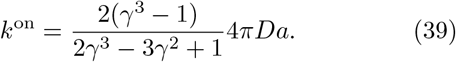

In the limit of *γ* → ∞ (random relocation of RNAPs in the whole system), the entering flux reduces to the classical result of diffusion-limited on-rate: *J* = 4*πDa*[*n*]_f,∞_ [63]. In our simulations, we take *γ* = 5, which leads to a *k*^on^ 1.4 times larger than that of the *γ* → ∞ case.

Finally, we define 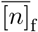 as the free RNAP concentration averaged over the space not occupied by binding sites after the system reaches the steady state. 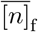 and [*n*]_f,∞_ are virtually the same because the binding sites only occupy a small fraction of the whole space (about 5%). Then the agent-based model can be mapped to the mean-field model by replacing [*n*]_f_ in Eq. (11) with the averaged free RNAP concentration 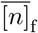.

### H. Agent-based simulation

Given the dissociation constants of the mean-field model, *K*_m_, *K*_r_, along with other parameters including *k*^on^ calculated in the above section and the initiation rates of rRNA and mRNA promoters (Table S1), we infer the values of 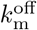 and 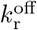 by 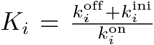 so that the mean-field model and the agent-based model are one-to-one correspondence. All the parameters in the simulation are for *E. coli* under the 30-min doubling time (Table S1), except for *K*_ns_ as we perform simulations under different effective volumes of nonspecific binding sites 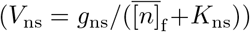. We tune *V*_ns_ by changing the offrate of nonspecific binding sites 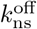 since 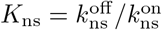 (Methods A). For the simulation of each *V*_ns_, we first esti-mate the total number of RNAPs given 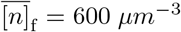 as the mean-field model according to Eq. (3).

In our simulation, the nucleoid is a cube; promoters and nonspecific binding sites are randomly distributed. Promoters and nonspecific binding sites are modeled as spheres with radius *a*. We model RNAPs as point particles, which are randomly distributed initially. We simulate the diffusion of RNAPs so that the x coordinate of an RNAP particle follows

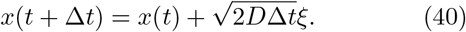

Here, *ξ* is a Gaussian random variable with zero mean and unit variance. The y and z coordinates follow similar dynamics. A free RNAP enters a binding site if the distance between them is smaller than *a* and the binding site is not occupied. We use the Gillespie Algorithm to determine the random dwell time and whether the RNAP should initiate transcription or leave after entering the binding site. An elongating RNAP becomes free after a fixed duration, which is the time for an RNAP to finish transcribing an operon, i.e., 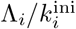 (Table S1).

We take the simulation time interval Δ*t* = 3 × 10^*−*7^ s and simulate for a total duration of *T* = 800 s. The detailed choice of Δ*t* and the corresponding error analysis are shown in Section A and Fig. S6 in the Supplementary Information. We calculate the free RNAP concentration distribution and RNA production rate using the data collected every second from *t*_1_ = 400 s to *T* = 800 s to ensure the system reaches a steady state. The free RNAP concentration distribution is averaged over each binding site of the same type and averaged over time. The RNA production rate is calculated as the total number of elongating RNAPs on rRNA (mRNA) genes divided by the time for an RNAP to finish transcribing, which is the same as the definition of the mean-field model (Eq. (5)). We repeat the simulations 20 times for each *V*_ns_ and calculate the mean and standard deviation of free RNAP concentration distribution across repeats. For each *V*_ns_, the correlation coefficient of mRNA and rRNA production rates is calculated across time by pooling all repeats together.

## ACKNOWLEDGMENTS

The research was funded by the National Key R&D Program of China (2021YFF1200500) and supported by grants from Peking-Tsinghua Center for Life Sciences. Figs. 1(a), 2(a-b), and 4(a-c) are created with BioRender.com.

## Notes

### Competing Interest Statement

The authors have declared no competing interest.

### Summary of Updates

The new section detailing the benefits and fitness costs of nonspecifically DNA-bound RNAPs has been added. Additionally, we have derived the partition rules for RNAPs, incorporating various types of sigma factors. All figures have been redrawn to significantly enhance readability.

