## Supplementary Information for "Buffering effects of nonspecifically DNA-bound RNA polymerases in bacteria"

Yichen Yan,[1] Tairan Li,[2] and Jie Lin[1,2]
[1] *Peking-Tsinghua Center for Life Sciences, Peking University, Beijing, China*
[2] *Center for Quantitative Biology, Peking University, Beijing, China*
(Dated: July 9, 2024)

TABLE S1. **Parameters for *E. coli* at doubling time 30 min for the mean-field model and agent-based simulations.** (1) The total number of RNAPs at doubling time 30 min is estimated to be 8000 [1, 2].
(2) At 30-min doubling time, there are two nucleoid lobes in a cell, segregated into two sublobes each. Each sublobe volume is 0.5 $\mu\text{m}^3$, so the total nucleoid volume is $V = 2\ \mu\text{m}^3$ [2].
(3) The free RNAP concentration is the number of free RNAPs divided by nucleoid volume: $[n]_\text{f} = n_\text{t} f_\text{free}/V$, where $f_\text{free} = 0.15$ (see Methods C in the main text).
(4-6) The number of rRNA operons ($g_\text{r}$) at 30-min doubling time is about 30 [1]. $g_\text{m}$ is estimated by dividing 4400 genes in a genome by three genes per promoter, and multiplied by three genomes per cell [1, 3, 4]. $g_\text{ns}$ is estimated as the genome size divided by the groove length of RNAP, which is 4.7 Mbp per genome multiplied by three genomes and divided by 16 bp.
(7-8) The maximum transcription initiation rates are the inverse of time interval between two initiations, which are about $1/0.6\ \text{s} = 1.7\ \text{s}^{-1}$ for rRNA genes and $1/2\ \text{s} = 0.5\ \text{s}^{-1}$ for mRNA genes [5, 6].
(9) At 30-min doubling time, the fraction of active RNAPs is measured to be 0.22 [1], and the fraction of specifically bound RNAPs is 0.54 [2]. Considering the promoter-bound RNAP number is small compared to the number of actively transcribing and pausing RNAPs [7], we calculate the pausing-active ratio for mRNA genes and rRNA genes together to be $(0.54 - 0.22)/0.22 = 1.5$. Here we assume the pausing-active ratio for rRNA genes and mRNA genes are the same ($\eta_\text{r} = \eta_\text{m} = 1.5$), and detailed assumptions do not affect our conclusions (Fig. S7).
(10-11) $\Lambda_r$ and $\Lambda_m$ are calculated by $\Lambda_i = k_i^\text{ini} \frac{L_i}{c_i}(1 + \eta_i)$, where $k_\text{m}^\text{ini} = 0.5\ \text{s}^{-1}$, $L_\text{m} = 3000$ nt, $c_\text{m} = 53$ nt/s; $k_\text{r}^\text{ini} = 1.7\ \text{s}^{-1}$, $L_\text{r} = 6500$ nt, $c_\text{r} = 85$ nt/s [1, 4, 6–8].
(12-14) $K_i$ ($K_\text{m}$ or $K_\text{r}$) is calculated by $n_\text{t} f_i = g_i(1 + \Lambda_i)\frac{[n]_\text{f}}{[n]_\text{f} + K_i}$; $K_\text{ns}$ is calculated by $n_\text{t} f_\text{ns} = g_\text{ns}\frac{[n]_\text{f}}{[n]_\text{f} + K_\text{ns}}$, where $f_\text{m} = 0.17$, $f_\text{r} = 0.38$, and $f_\text{ns} = 0.30$ (Methods C in the main text). The estimate of $K_\text{ns}$ is close to previous measurement results [9, 10], but smaller than the estimate in [7] where the nonspecific RNAP fraction is overestimated.
(15) $D = 0.7\ \mu\text{m}^2\text{s}^{-1}$ is the diffusion constant of free RNAPs measured at 30-min doubling time [2].
(16) The radius of binding sites is set to be slightly larger than the width of double-stranded DNA [11].
(17) An RNAP can slide up to 1000 bp along DNA before detachment [12], which means it can scan a cylindrical (radius $= a$) volume of $10^{-5}\mu\text{m}^{-3}$. In our simulation we use a sphere to represent this volume, which leads to a radius of about 5 times $a$.

| **Parameters** | **Symbols** | **Values** |
|---|---|---|
| (1) Total RNAP number | $n_\text{t}$ | 8000 |
| (2) Nucleoid volume | $V$ | $2\ \mu\text{m}^3$ |
| (3) Free RNAP concentration | $[n]_\text{f}$ | $600\ \mu\text{m}^{-3}$ |
| (4) Number of rRNA promoters | $g_\text{r}$ | 30 |
| (5) Number of mRNA promoters | $g_\text{m}$ | 4400 |
| (6) Number of nonspecific binding sites | $g_\text{ns}$ | $9 \times 10^5$ |
| (7) maximum transcription initiation rate of rRNA genes | $k_\text{r}^\text{ini}$ | $1.7\ \text{s}^{-1}$ |
| (8) maximum transcription initiation rate of mRNA genes | $k_\text{m}^\text{ini}$ | $0.5\ \text{s}^{-1}$ |
| (9) Pausing-active ratio for rRNA genes and mRNA genes | $\eta_\text{r}$, $\eta_\text{m}$ | 1.5 |
| (10) Maximum capacity for elongating RNAPs per active rRNA operon | $\Lambda_\text{r}$ | 325 |
| (11) Maximum capacity for elongating RNAPs per active mRNA operon | $\Lambda_\text{m}$ | 75 |
| (12) Dissociation constant of rRNA promoters | $K_\text{r}$ | $1.3 \times 10^3\ \mu\text{m}^{-3}$ |
| (13) Dissociation constant of mRNA promoters | $K_\text{m}$ | $1.4 \times 10^5\ \mu\text{m}^{-3}$ |
| (14) Dissociation constant for nonspecific binding | $K_\text{ns}$ | $2.3 \times 10^5\ \mu\text{m}^{-3}$ |
| (15) Diffusion constant of free RNAPs | $D$ | $0.7\ \mu\text{m}^2\text{s}^{-1}$ |
| (16) Radius of binding sites | $a$ | $3 \times 10^{-3}\ \mu\text{m}$ |
| (17) Relocation distance parameter | $\gamma$ | 5 |

TABLE S2. **Parameter ranges of the model considering sigma and anti-sigma factors.** All genes are divided into three types as mentioned in the main text (genes A and B recognized by $\sigma^1$, and genes C recognized by $\sigma^2$). Each parameter is randomly picked from a $log_{10}$-scale uniform distribution within its range. We use Matlab 2023a 'fsolve' function to numerically solve Eqs. (22-23,25-28) in the main text, and discard the parameter sets that fail to give solutions.
(1) The number of housekeeping sigma factors $\sigma^{70}$ is measured to be 500 to 20000 even when the total RNAP number is not much changed [13–17]. Other alternative sigma factors like $\sigma^E, \sigma^H, \sigma^S$ are about a few hundreds [13–18]. Therefore, the concentration ranges of $[\sigma^1]_\mathrm{t}$ and $[\sigma^2]_\mathrm{t}$ are both set to be [250-10000] given a nucleoid volume of 2 $\mu$m$^3$. If $[\sigma^1]_\mathrm{t} > [\sigma^2]_\mathrm{t}$, it may correspond to the case where genes A and B are controlled by $\sigma^{70}$, and genes C are controlled by $\sigma^{Alt}$.
(2) The number of anti-sigma factors is about the same magnitude as sigma factors [14], so we set their ranges to be the same.
(3) For both $E\sigma^{70}$ and $E\sigma^S$, the backward rates are measured to be on the order of $10^{-3}$ s$^{-1}$ [19, 20]. Here we scan two orders of magnitude with the experimental results included in the middle.
(4) The dissociation constants of holoenzyme can vary from 0.2 to 2000 $\mu$m$^3$ for any kind of holoenzymes, and most are on the order of tens [17, 19, 21–23]. Here we still scan two orders of magnitude from $10^0 - 10^2$ $\mu$m$^3$.
(5) The dissociation constants of $\sigma$·anti-$\sigma$ complex are about the order of tens [24]. Again we set the range to be $10^0 - 10^2$ $\mu$m$^3$.
(6) By comparing the number of RNAPs on genes $i(j)$ to the number of free RNAPs, we get the folloing relationship between the RNAP fractions and effective volumes: $\frac{f_{i(j)}}{f_\mathrm{free}} = \frac{n_{i(j)}}{n_\mathrm{f}} = \frac{[E\sigma^j]_\mathrm{f} V_{i(j)}}{[n]_\mathrm{f} V} = \frac{\beta_j}{1+\sum_j \beta_j}\frac{V_{i(j)}}{V}$. As discussed in Methods C regarding the growth-rate-dependent RNAP partition fractions, the fraction ratio $\frac{f_{i(j)}}{f_\mathrm{free}}$ should be on the scale of 1. Considering $\frac{\beta_j}{1+\sum_j \beta_j}$ is smaller than 1 and $V = 2$ $\mu$m$^3$, we scan the effective volume on the range of 1-10.
(7) The maximum elongating capacity we estimate in Table S1 is on the order of tens to a few hundreds. Here considering some variations in the maximum initiation rates and gene length, we scan a tenfold range with $\Lambda_\mathrm{m}$ value in Table S1 logarithmically in the middle.

| **Parameters** | **Symbols** | **Ranges** | **Units** |
|---|---|---|---|
| (1) Total $\sigma$ concentration | $[\sigma^1]_\mathrm{t}, [\sigma^2]_\mathrm{t}$ | [250,10000] | $\mu$m$^{-3}$ |
| (2) Total anti-$\sigma$ factor concentration | $[Anti^1]_\mathrm{t}, [Anti^2]_\mathrm{t}$ | [250,10000] | $\mu$m$^{-3}$ |
| (3) Backward rate of holoenzyme formation | $k^\mathrm{b}_{E\sigma^1}, k^\mathrm{b}_{E\sigma^2}$ | $[10^{-4}, 10^{-2}]$ | s$^{-1}$ |
| (4) Dissociation constant of holoenzyme | $K_{E\sigma^1}, K_{E\sigma^2}$ | [1,100] | $\mu$m$^{-3}$ |
| (5) Dissociation constant of $\sigma$·anti-$\sigma$ complex | $K_{\sigma^1 \cdot Anti^1}, K_{\sigma^2 \cdot Anti^2}$ | [1,100] | $\mu$m$^{-3}$ |
| (6) Effective volume $V_{i(j)}$ | $V_{\mathrm{A}(1)}, V_{\mathrm{B}(1)}, V_{\mathrm{C}(2)}$ | [1,10] | $\mu$m$^3$ |
| (7) Maximum capacity for elongating RNAPs $\Lambda_{i(j)}$ | $\Lambda_{\mathrm{A}(1)}, \Lambda_{\mathrm{B}(1)}, \Lambda_{\mathrm{C}(2)}$ | $[25 - 250]$ | 1 |

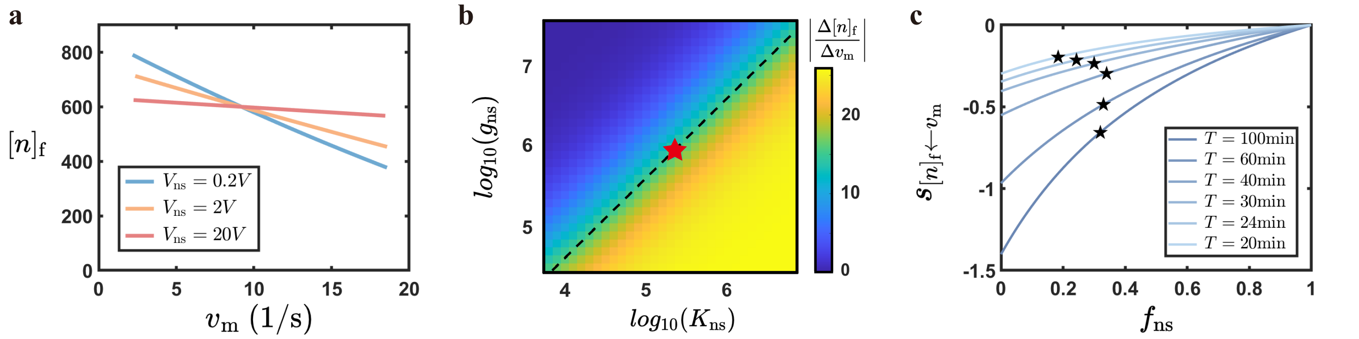


FIG. S1. **The free RNAP concentration is buffered by nonspecific RNAPs against sudden changes in mRNA expression.** (a) The free RNAP concentration vs. the mRNA production rates under different effective volumes of nonspecific binding. The free RNAP concentration before perturbation under different $V_\mathrm{ns}$ is fixed at $[n]_\mathrm{f} = 600\ \mu\mathrm{m}^{-3}$ (Table S1). The $V_\mathrm{ns} = 2V$ line corresponds to the parameters of *E. coli*. (b) The absolute slopes of $[n]_\mathrm{f}$ over $v_\mathrm{m}$ vs. $K_\mathrm{ns}$ and $g_\mathrm{ns}$. The slopes are equal as long as $g_\mathrm{ns}/K_\mathrm{ns}$ is the same (indicated by the dashed line with slope 1 in the logarithmic coordinates). The red star marks the corresponding values of *E. coli* estimated under 30-min doubling time (Table S1). (c) The sensitivity of free RNAP concentration to the changes in mRNA production rates (Eq. (15) in the main text) as a function of the nonspecific RNAP fraction under different growth rates. The stars mark the $f_\mathrm{ns}$ values of *E. coli* and the corresponding y-values.

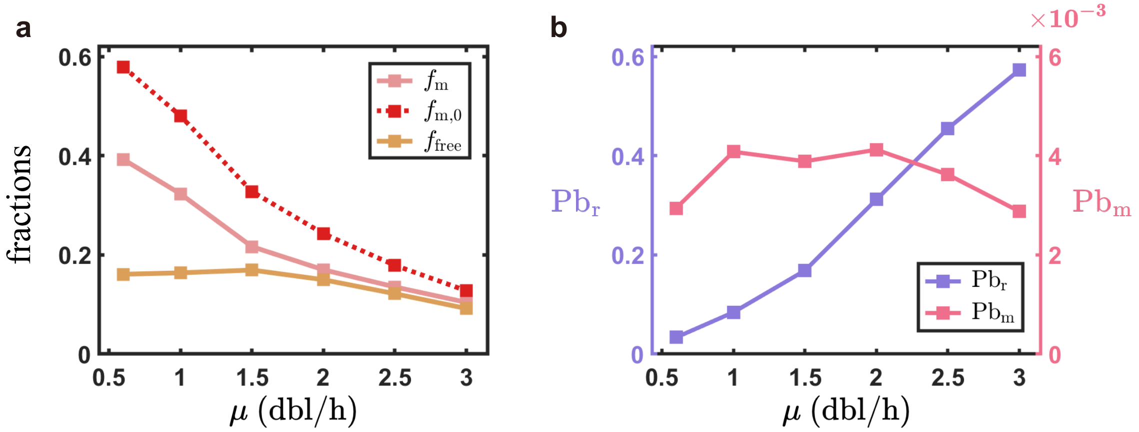


FIG. S2. **Growth-rate-dependent RNAP partition fractions and RNAP binding probabilities.** (a) Fractions of free RNAPs ($f_{\mathrm{free}}$), RNAPs on mRNA genes ($f_{\mathrm{m}}$), and RNAPs on mRNA genes if nonspecific binding is absent ($f_{\mathrm{m},0}$) across different growth rates ($\mu$). (b) Probabilities of promoter bound by an RNAP of rRNA genes ($\mathrm{Pb_r}$) and mRNA genes ($\mathrm{Pb_m}$). The growth -rate-dependent $f_{\mathrm{r}}$, $f_{\mathrm{r},0}$ and $f_{\mathrm{ns}}$ are shown in Fig. 1(d) in the main text.

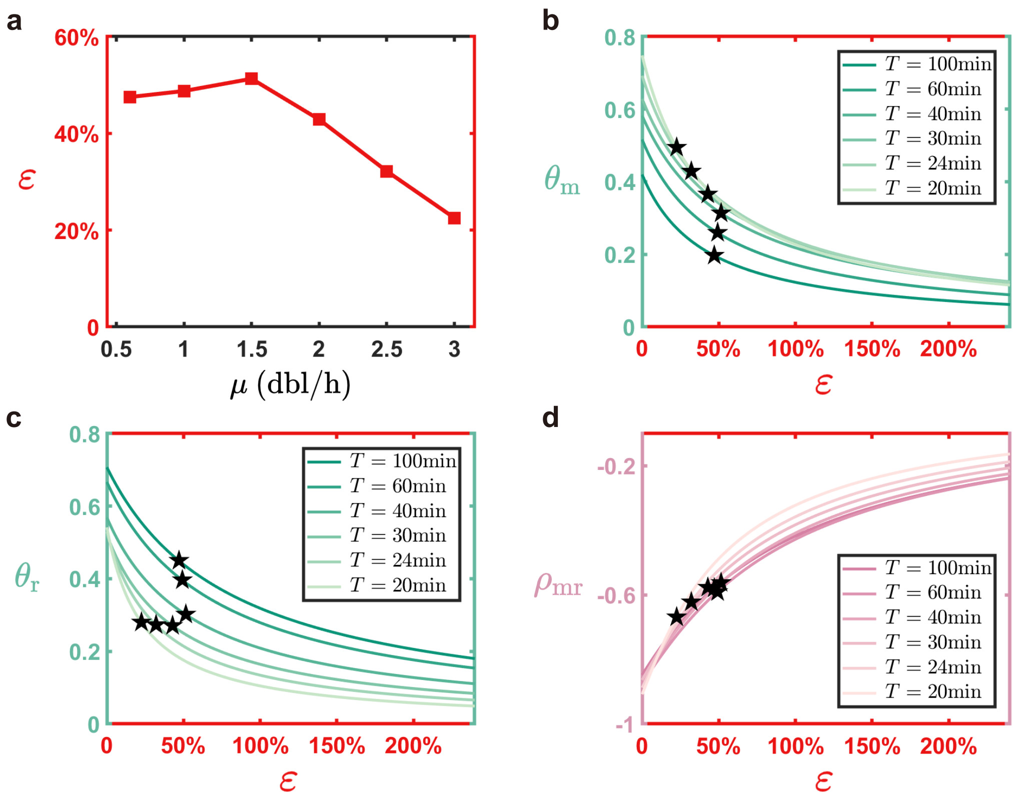


FIG. S3. **The benefit of nonspecific RNAPs compared to the cost measured by the excess factor $\varepsilon$.** (a) The cost of nonspecific RNAPs under different growth rates, quantified by the excess factor $\varepsilon$ ($f_{\mathrm{ns}}/(1-f_{\mathrm{ns}})$). (b-c) The crosstalk factors for mRNA genes (b) and rRNA genes (c) as a function of the excess factor across six different growth rates. Here, $T$ is the doubling time, the inverse of the growth rate. The stars mark the corresponding values of *E. coli*. (d) The correlation coefficient of mRNA and rRNA production rates (Eq. (21) in the main text) as a function of the excess factor. The stars mark the corresponding values of *E. coli*. Here $D_{\mathrm{m}} = D_{\mathrm{r}} = 0.2$, and using different fluctuation degrees does not affect our main conclusion (Fig. S4).

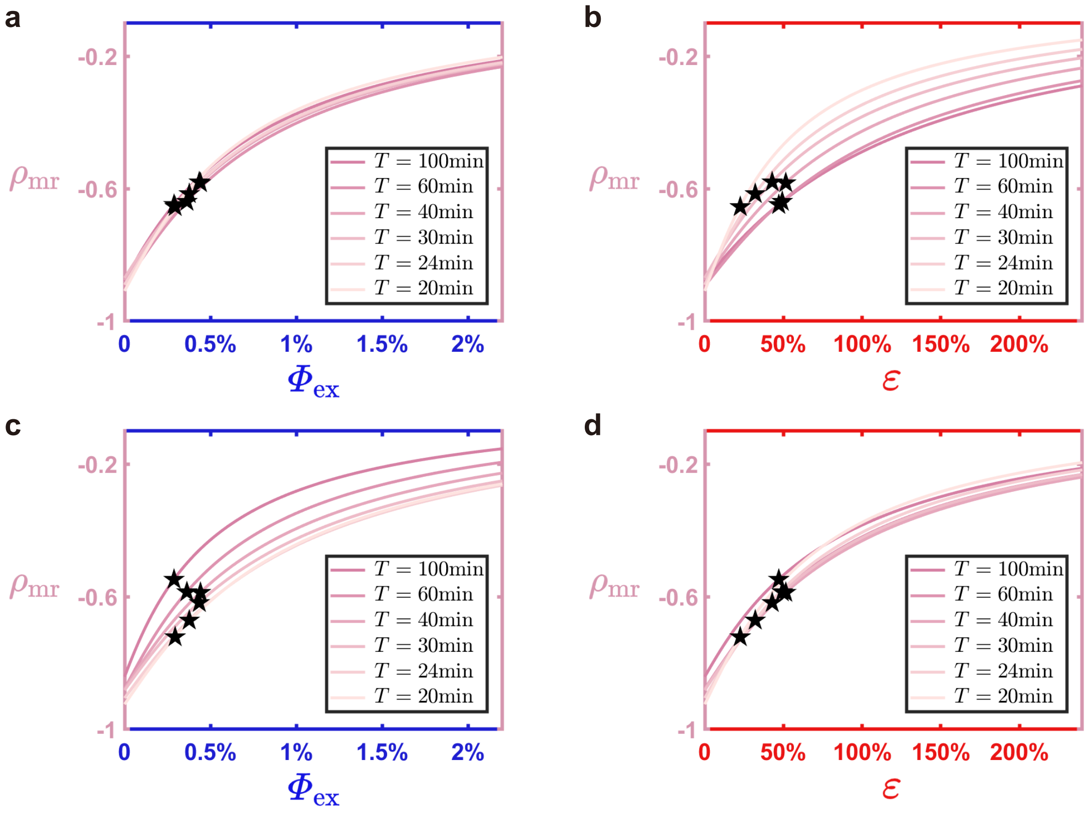


FIG. S4. **Nonspecific RNAPs attenuate gene expression crosstalk regardless of the mRNA and rRNA noise levels.** (a-b) The correlation coefficient of mRNA and rRNA production rates as a function of the proteome fraction of excess nonspecific RNAPs (a) or the excess factor (b) when $D_{\mathrm{m}} = 2D_{\mathrm{r}}$. Here, $T$ is the doubling time, the inverse of the growth rate. The stars mark the corresponding values of *E. coli.* (c-d) Same as (a-b) except $D_{\mathrm{r}} = 2D_{\mathrm{m}}$. Comparing these results to Fig. 2(d) and Fig. S3(d) where we set $D_{\mathrm{m}} = D_{\mathrm{r}}$, we conclude that the attenuation effect of nonspecific RNAPs on the gene expression crosstalk is not affected by the noise levels of mRNA and rRNA gene expression. Here, the absolute values of $D_{\mathrm{m}}$ and $D_{\mathrm{r}}$ are not important because the correlation factor is not changed as long as $D_{\mathrm{m}}/D_{\mathrm{r}}$ keeps the same (Eq. (21) in the main text).

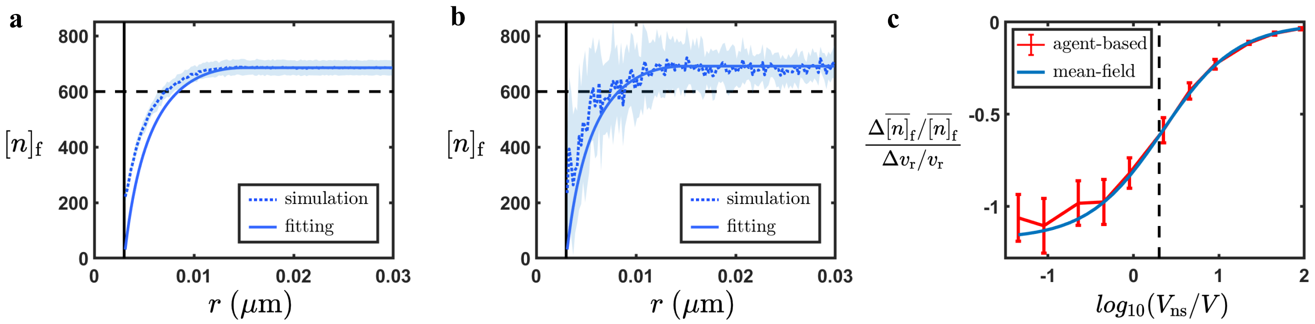


FIG. S5. **The agent-based simulations agree well with the theoretical free RNAP distribution (a-b) and mean-field predictions regarding the buffering effects of nonspecific RNAPs (c).** (a-b) The free RNAP concentration distribution around a nonspecific binding site (a) and a promoter (b) in agent-based simulations (dotted curve) can be well fitted by the analytical predictions (Eqs. (33, 35) in the main text, solid curve). Nevertheless, the free RNAP concentration far from a binding site slightly exceeds the preassigned value ($[n]_{\mathrm{f},\infty} = 600\ \mu\mathrm{m}^3$, dashed line), and there is a finite free RNAP concentration at the boundary of binding sites ($r = a$, vertical solid line). The shaded areas represent the standard deviation from 20 repeats. For each repeat, the free RNAP concentration is calculated every $6 \times 10^{-4}\ \mu$m from $r = a$ and is averaged over all binding sites of the same type. The free RNAP concentration around a promoter fluctuates more than a nonspecific binding site due to the lower copy number of promoters. (c) The sensitivity of the averaged free RNAP concentration to the rRNA production rate ($\frac{\Delta\overline{[n]}_\mathrm{f}/\overline{[n]}_\mathrm{f}}{\Delta v_\mathrm{r}/v_\mathrm{r}}$) upon a sudden rRNA gene up-regulation *vs.* the effective volume of nonspecific binding sites $V_\mathrm{ns}$. We up-regulate the expression of rRNA genes by setting $K_\mathrm{r,up} = 0.2K_\mathrm{r}$, specifically by changing $k_\mathrm{r}^\mathrm{off}$. This change is performed at $T = 800$ s, and we run another 800 s to ensure the system reaches the new steady state. The free RNAP concentrations before and after regulation are both averaged over their last 400 s. The blue curve is the mean-field prediction by Eq. (3) in the main text, and the red curve is the result of the agent-based simulations. The dashed line represents the estimated parameters of *E. coli*. In this figure, we use the parameters in Table S1 unless otherwise mentioned. We change $V_\mathrm{ns}$ by changing the dissociation rate $K_\mathrm{ns}$, specifically by changing the off-rate $k_\mathrm{ns}^\mathrm{off}$. Data are shown as mean $\pm$ s.d. (20 repeats)

.

#### A. The free RNAP concentration distribution and error analysis

When the time interval in simulation is not small enough, the average length of each random walk ($\sqrt{6D\Delta t}$) will be comparable to or even larger than the sizes of the binding sites. Therefore, it is possible that a particle is outside the binding site at the start and the end of the time interval, but during the time interval, it actually enters the binding site. This error due to a finite time interval results in a smaller flux entering the binding sites than the theoretical prediction and a finite free RNAP concentration at the surface of binding sites (Figs. S5(a-b) and S6(b-c)). Intuitively, an underestimation of bound RNAPs leads to an overestimation of free RNAPs. To be specific, the simulated on-rate $k_{\rm on}^{\rm sim}$ is smaller than the expected value given by Eq. (39) in the main text due to the underestimation. According to $K_i = \frac{k_i^{\rm off}+k_i^{\rm ini}}{k_i^{\rm on}}$, smaller $k_{\rm sim}^{\rm on}$ leads to larger dissociation constants, indicating lower binding affinities. Therefore, the free RNAP concentration becomes higher than expected. As $\Delta t$ decreases, the distributions approach the theoretical predictions (Fig. S6(b, c)).

To test our idea explicitly, we calculate $k_{\rm sim}^{\rm on}$ and the free RNAP concentration distributions under different $\Delta t$. For simplicity, we simulate a single binding site with radius $a$ and generate random particles following the free RNAP concentration distributions according to Eqs. (33,35) in the main text ($\gamma = 5$ as we use in all simulations). Here, we take an unrealistically large $[n]_{\rm f,\infty} = 10^{10}$ $\mu$m$^{-3}$ to minimize noise. We remark that the purpose of the simulation is to demonstrate the error due to the finite time interval $\Delta t$ and the value of $[n]_{\rm f,\infty}$ does not affect our conclusion as it is canceled out in the calculation. We let RNAPs perform random walk with a time interval $\Delta t$. If an RNAP is located inside the binding site at the end of each time interval, we relocate it randomly inside a shell with inner diameter $a$ and outer diameter $\gamma \times a$ as we do in the main text.

After a duration of $T = 10^{-5}$ s, we count the total number of absorbed particles, $N$. The simulated on-rate is calculated as $k_{\rm sim}^{\rm on} = \frac{N}{T[n]_{\rm f,\infty}}$. Since $N$ is proportional to $[n]_{\rm f,\infty}$ because different particles are independent, the results do not depend on the value of $[n]_{\rm f,\infty}$. We calculate the ratio of $k_{\rm sim}^{\rm on}$ to the theoretical expectation $k_{\rm theory}^{\rm on}$ and find that a finite time interval makes the simulated $k_{\rm sim}^{\rm on}$ smaller than the expectation (Fig. S6(a)). To balance the accuracy and simulation time, we choose $\Delta t = 3 \times 10^{-7}$ s, which already provides good results (Figs. 4(d-e) and Fig. S5).

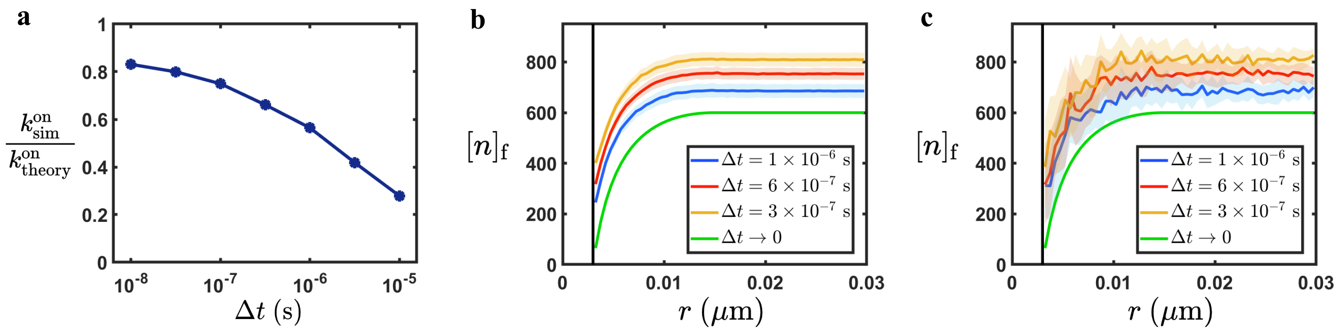


FIG. S6. **The time interval $\Delta t$ influences the simulated on rate $k_{\rm sim}^{\rm on}$ and the free RNAP concentration.** (a) The ratio of the simulated on rate $k_{\rm sim}^{\rm on}$ from the simplified simulation with a single binding site to the theoretical prediction, Eq. (39) in the main text. (b-c) The free RNAP concentration distribution around a nonspecific binding site (b) and a promoter (c) in agent-based simulations under different $\Delta t$. The $\Delta t \to 0$ lines are the theoretical predictions. The solid line marks the boundary of binding sites ($r = a$). All the results here are before the regulation of rRNA genes. The shaded areas represent the standard deviation of 20 repeats. Here for each repeat, the free RNAP concentration is calculated every $1.5 \times 10^{-3}$ $\mu$m from $r = a$ and is averaged over all binding sites of the same type. Data are shown as mean ± s.d.

### B. Our conclusion is robust regardless of detailed assumptions about the pausing-active ratio $\eta$

In the main text, we assume the pausing-active ratios for mRNA genes and rRNA genes are the same, which are 1.5 at 30-min doubling time, decrease with growth rate and approach 0 when the growth rate is 5 $h^{-1}$ (i.e., doubling time $T = 12$ min). The corresponding results of growth-rate-dependent RNAP fractions and binding probabilities are presented in Fig. 1(c) and Fig. S2, which apply to Figs. 1(d-e), 2, and S3. We also consider other reasonable assumptions about $\eta_{\mathrm{m}}$ and $\eta_{\mathrm{r}}$. The ppGpp mainly induces the pausing during mRNA synthesis, while rRNA synthesis is less affected [25]. Therefore, we can assume the pausing-active ratio of rRNA genes is smaller than that of mRNA genes, and is constant across different growth rates. This leads to the following estimation: $\eta_{\mathrm{m}} = 3.6 - 0.5\mu$; $\eta_{\mathrm{r}} = 1.0$. Fig. S7(a-b) shows the buffering effects of nonspecific RNAPs on free RNAP concentration upon sudden changes in gene expression using the new set of parameters, corresponding to Fig. 1(d-e) in the main text. Fig. S7(c-i) shows the cost and benefit of nonspecific RNAPs, corresponding to Fig. 2 in the main text and Fig. S3. In summary, we find the detailed choices of the pausing-active ratios do not affect our main conclusions.

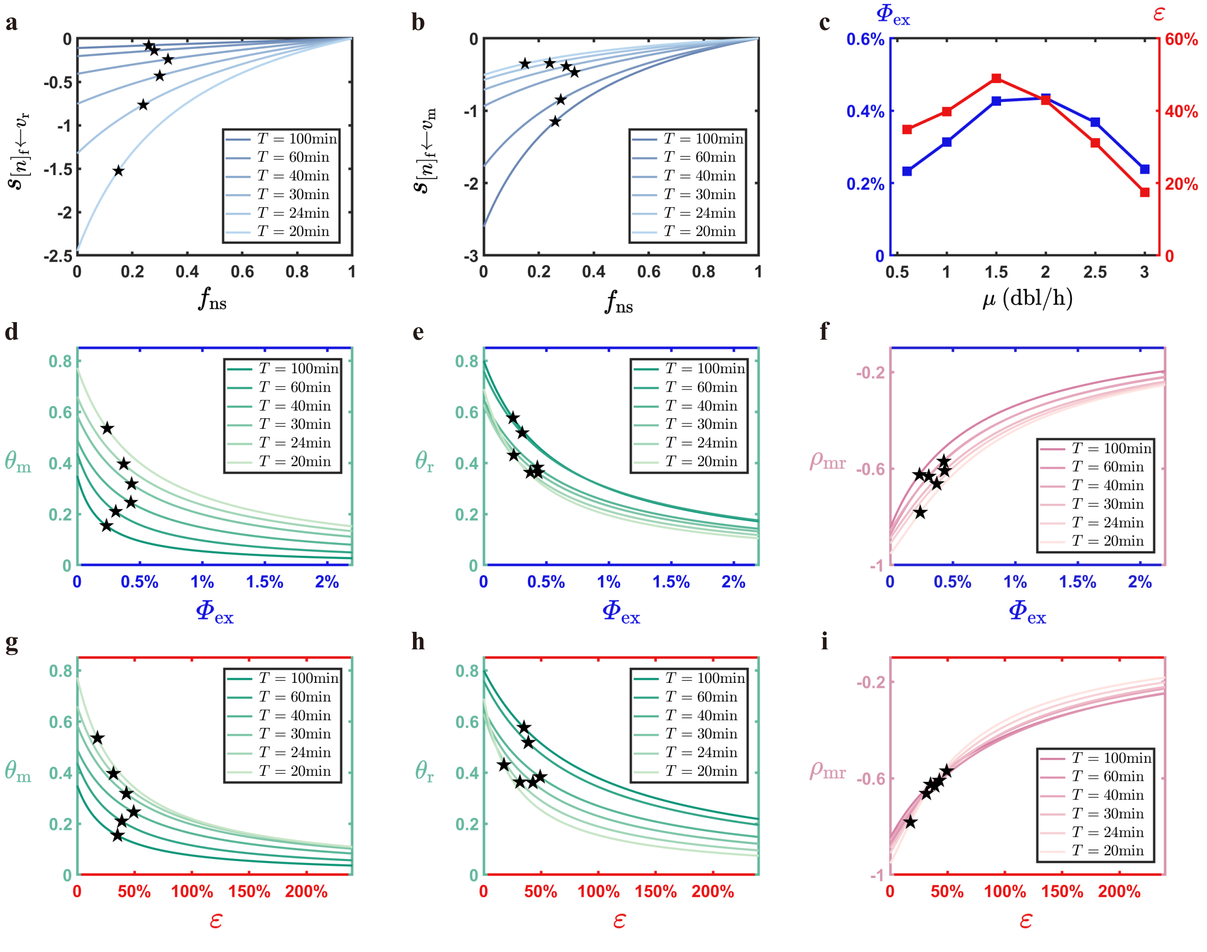


FIG. S7. **Results of the mean-field model using another set of pausing-active ratio parameters.** (a-b) The sensitivity of free RNAP concentration to the changes of rRNA (a) and mRNA (b) production rates as a function of the nonspecific RNAP fraction under different growth rates. Here, $T$ is the doubling time, the inverse of the growth rate, which also applies to other panels. The stars mark the $f_{\mathrm{ns}}$ of *E. coli* and the corresponding y-values. (c) The two indexes of cost of nonspecific RNAPs under different growth rates. The blue color marks the proteome fraction of excess nonspecific RNAPs ($\Phi_{\mathrm{ex}}$), and the red color marks the excess factor $\varepsilon$, and . (d-f) The benefit of nonspecific RNAPs *vs* the proteome fraction of excess nonspecific RNAPs ($\Phi_{\mathrm{ex}}$, blue). Nonspecific RNAPs promote the gene expression accuracy of mRNA genes (d) and rRNA genes (e) and attenuate the gene expression crosstalk between mRNA and rRNA genes (f) at a low cost. The stars mark the corresponding values of *E. coli*. (g-i) Same as (d-f), but the x-axis is changed to the excess factor ($\varepsilon$, red).
